# Polygenic adaptation and convergent evolution across both growth and cardiac genetic pathways in African and Asian rainforest hunter-gatherers

**DOI:** 10.1101/300574

**Authors:** Christina M. Bergey, Marie Lopez, Genelle F. Harrison, Etienne Patin, Jacob Cohen, Lluis Quintana-Murci, Luis B. Barreiro, George H. Perry

**Affiliations:** Department of Anthropology, Pennsylvania State University, University Park, Pennsylvania, U.S.A.; Department of Biology, Pennsylvania State University, University Park, Pennsylvania, U.S.A.; Unit of Human Evolutionary Genetics, Institut Pasteur, Paris, France.; Centre National de la Recherche Scientifique UMR 2000, Paris, France.; Center of Bioinformatics, Biostatistics and Integrative Biology, Institut Pasteur, Paris, France.; Université de Montréal, Centre de Recherche CHU Sainte-Justine, Montréal, Canada.; Huck Institutes of the Life Sciences, Pennsylvania State University, University Park, Pennsylvania, U.S.A.

## Abstract

Different human populations facing similar environmental challenges have sometimes evolved convergent biological adaptations, for example hypoxia resistance at high altitudes and depigmented skin in northern latitudes on separate continents. The pygmy phenotype (small adult body size), a characteristic of hunter-gatherer populations inhabiting both African and Asian tropical rainforests, is often highlighted as another case of convergent adaptation in humans. However, the degree to which phenotypic convergence in this polygenic trait is due to convergent vs. population-specific genetic changes is unknown. To address this question, we analyzed high-coverage sequence data from the protein-coding portion of the genomes (exomes) of two pairs of populations, Batwa rainforest hunter-gatherers and neighboring Bakiga agriculturalists from Uganda, and Andamanese rainforest hunter-gatherers (Jarawa and Onge) and Brahmin agriculturalists from India. We observed signatures of convergent positive selection between the Batwa and Andamanese rainforest hunter-gatherers across the set of genes with annotated ‘growth factor binding’ functions (*p* < 0.001). Unexpectedly, for the rainforest groups we also observed convergent and population-specific signatures of positive selection in pathways related to cardiac development (e.g. ‘cardiac muscle tissue development’; *p* = 0.001). We hypothesize that the growth hormone sub-responsiveness likely underlying the pygmy phenotype may have led to compensatory changes in cardiac pathways, in which this hormone also plays an essential role. Importantly, in the agriculturalist populations we did not observe similar patterns of positive selection on sets of genes associated with either growth or cardiac development, indicating that our results most likely reflect a history of convergent adaptation to the similar ecology of rainforest hunter-gatherers rather than a more common or general evolutionary pattern for human populations.

## Introduction

Similar ecological challenges may repeatedly result in similar evolutionary outcomes, and many instances of phenotypic convergence arising from parallel changes in the same genetic loci have been uncovered (reviewed in [1–3]). Many examples of convergent genetic evolution reported to date are for simple monogenic traits, for example depigmentation in independent populations of Mexican cave fish living in lightless habitats [4, 5] and persistence of the ability to digest lactose in adulthood in both European and African agriculturalist/pastoralist humans [6]. Most biological traits, however, are highly polygenic. Since the reliable detection of positive selection in aggregate on multiple loci of individually small effect (i.e., polygenic adaptation) is relatively difficult [7–11], the extent to which convergent genetic changes at the same loci and functional pathways or changes affecting distinct genetic pathways may underlie these complex traits is less clear.

Human height is a classic example of a polygenic trait with approximately 800 known loci significantly associated with stature in Europeans collectively accounting for 27.4% of the heritable portion of height variation in this population [12]. A stature phenotype also represents one of most striking examples of convergent evolution in humans. Small body size (or the “pygmy” phenotype, e.g. average adult male stature <155 cm) appears to have evolved independently in rainforest hunter-gatherer populations from Africa, Asia, and South America [13], as groups on different continents do not share common ancestry to the exclusion of nearby agriculturalists [14, 15]. Positive correlations between stature and the degree of admixture with neighboring agriculturalists have confirmed that the pygmy phenotype is, at least in part, genetically mediated and therefore potentially subject to natural selection [16–20].

Indeed, previous population genetic studies have identified signatures of strong positive natural selection across the genomes of various worldwide rainforest hunter-gatherer groups [15, 19, 21, 22]. In some cases, the candidate positive selection regions were significantly enriched for genes involved in growth processes and pathways [15, 19]. However, in one rainforest hunter-gatherer population, the Batwa from Uganda, an admixture mapping approach was used to identify 16 genetic loci specifically associated with the pygmy phenotype [17]. While these genomic regions were enriched for genes involved in the growth hormone pathway and for variants associated with stature in Europeans, there was no significant overlap between the pygmy phenotype-associated regions and the strongest signals of positive selection in the Batwa genome. Rather, subtle shifts in allele frequencies were observed across these regions in aggregate, consistent with a history of polygenic adaptation for the Batwa pygmy phenotype [17] and underscoring the importance of using different types of population genetic approaches to study the evolutionary history of this trait. Similar studies focused on other rainforest hunter-gatherer groups have found enrichment for signatures of selection on genes involved in growth [15] and various growth factor signaling pathways [19], immunity [19, 21, 22], metabolism [19, 21, 22], development [15, 22], and reproduction [19, 21, 22].

Here, we investigate population-specific and convergent patterns of positive selection in African and Asian hunter-gatherer populations using genome-wide sequence data from two sets of populations: the Batwa rainforest hunter-gatherers of Uganda in East Africa and the nearby Bakiga agriculturalists [23], and the Jarawa and Onge rainforest hunter-gatherers of the Andaman Islands in South Asia and the Uttar Pradesh Brahmin agriculturalists from mainland India [24, 25]. We specifically test whether convergent or population-specific signatures of positive selection, as detected both with ‘outlier’ tests designed to identify strong signatures of positive selection and tests designed to identify signatures of polygenic adaptation, are enriched for genes with growth-related functions. After studying patterns of convergent-and population-specific evolution in the Batwa and Andamanese hunter-gatherers, we then repeat these analyses in the paired Bakiga and Brahmin agriculturalists to evaluate whether the evolutionary patterns most likely relate to adaptation to hunter-gatherer subsistence in rainforest habitats, rather than being more generalized evolutionary patterns for human populations.

## Results

We sequenced the protein coding portions of the genomes (exomes) of 50 Batwa rainforest hunter-gatherers and 50 Bakiga agriculturalists (dataset originally reported in [23]), identified single nucleotide polymorphisms (SNPs), and analyzed the resultant data alongside those derived from published whole genome sequence data for 10 Andamanese rainforest hunter-gatherers and 10 Brahmin agriculturalists (dataset from [25]). We restricted our analysis to exonic SNPs, for comparable analysis of the Asian whole genome sequence data with the African exome sequence data. To polarize allele frequency differences observed between each pair of hunter-gatherer and agriculturalist populations, we merged these data with those from outgroup comparison populations from the 1000 Genomes Project [26]: exome sequences of 30 unrelated British individuals from England and Scotland (GBR) for comparison with the Batwa/Bakiga data, and exome sequences of 30 Luhya individuals from Webuye, Kenya (LWK) for comparison with the Andamanese/Brahmin data. Outgroup populations were selected for genetic equidistance from the test populations. While minor levels of introgression from a population with European have been observed for the Batwa and Bakiga [23, 27], PBS is relatively robust to low levels of admixture [28].

To identify regions of the genome that may have been affected by positive selection in each of our test populations, we computed the population branch statistic (PBS; [29]) for each exonic SNP identified among or between the Batwa and Bakiga, and Andamanese and Brahmin populations (Fig. S1, S2; Table S15). PBS is an estimate of the magnitude of allele frequency change that occurred along each population lineage following divergence of the most closely related populations, with the allele frequency information from the outgroup population used to polarize frequency changes to one or both branches. Larger PBS values for a population reflect greater allele frequency change on that branch, which in some cases could reflect a history of positive selection [29].

For each analyzed population, we computed a PBS selection index for each gene by comparing the mean PBS for all SNPs located within that gene to a distribution of values estimated by shuffling SNP-gene associations (without replacement) and re-computing the mean PBS value for that gene 100,000 times (Table S17). The PBS selection index is the percentage of permuted values that is higher than the actual (observed) mean PBS value for that gene. Per-gene PBS selection index values were not significantly correlated with gene size (linear regression of log adjusted selection indices against gene length: adjusted *R*^2^ = −2.74 × 10^−5^, *F*-statistic *p* = 0.81; Fig. S3), suggesting that this metric is not overtly biased by gene size.

Convergent evolution can operate at different scales, including on the same mutation or amino acid change, different genetic variants between populations but within the same genes, or across a set of genes involved in the same biomolecular pathway or functional annotation. Given that our motivating phenotype is a complex trait and signatures of polygenic adaptation are expected to be relatively subtle and especially difficult to detect at the individual mutation and gene levels, in this study we principally consider patterns of convergence versus population specificity at the functional pathway/annotation level. We do note that when we applied the same approaches described in this study to individual SNPs, we identified several individual alleles with patterns of convergent allele frequency evolution between the Batwa and Andamanese that may warrant further study (Table S16), including a nonsynonymous SNP in the gene *FIG4*, which when disrupted in mice results in a phenotype of small but proportional body size [30]. However, likely related to the above-discussed challenges of identifying signatures of polygenic adaptation at the locus-specific level, the results of our individual SNP and gene analyses were otherwise largely unremarkable, and thus the remainder of our report and discussion focuses on pathway-level analyses.

### Outlier signatures of strong convergent and population-specific selection

The set of genes with the lowest (outlier) PBS index values for each population may be enriched for genes with histories of relatively strong positive natural selection. We used a permutation-based analysis to test whether curated sets of genome-wide growth-associated genes (4 lists tested separately ranging from 266-3,996 genes; 4,888 total genes; Suppl. Text) or individual Gene Ontology (GO) annotated functional categories of genes (GO categories with fewer than 50 genes were excluded) have significant convergent excesses of genes with low PBS selection index values (< 0.01) in both of two cross-continental populations, for example the Batwa and Andamanese. Specifically, we first used Fisher’s exact tests to estimate the probability that the number of genes with PBS selection index values < 0.01 was greater than that expected by chance, for each functional category set of genes and population. We then reshuffled the PBS selection indices across all genes 1,000 different times for each population to generate distributions of permuted enrichment p-values for each functional category set of genes. We compared our observed Batwa and Andamanese Fisher’s exact test p-values to those from the randomly generated distributions as follows. We computed the joint probability of the null hypotheses for both the Andamanese and Batwa being false as (1 − *p*_*Batwa*_)(1 − *p*_*Andamanese*_) where *p*_*Batwa*_ and *p*_*Andamanese*_ are the p-values of the Fisher’s exact test, and we compared this joint probability estimate to the same statistic computed for the p-values from the random iterations. We then defined the p-value of our empirical test for convergent evolution as the probability that this statistic was more extreme (lower) for the observed values than for the randomly generated values. The resultant p-value summarizes the test of the null hypothesis that both results could have been jointly generated under random chance. While each individual population’s outlier-based test results are not significant after multiple test correction, this joint approach provides increased power to identify potential signatures of convergent selection by assessing the probability of obtaining two false positives in these independent samples.

Several GO biological processes were significantly overrepresented—even when accounting for the number of tests performed—among the sets of genes with outlier signatures of positive selection in both the Batwa and Andamanese hunter-gatherer populations (empirical test for convergence *p* < 0.005; Table S1; Fig. 1A). These GO categories include ‘limb morphogenesis’ (GO:0035108; empirical test for convergence *p* < 0.001; *q* < 0.001; Batwa: genes observed = 5, expected = 1.69, Fisher’s exact *p* = 0.027; Andamanese: observed = 6, expected = 2.27, Fisher’s exact *p* = 0.025).

**Figure 1:**
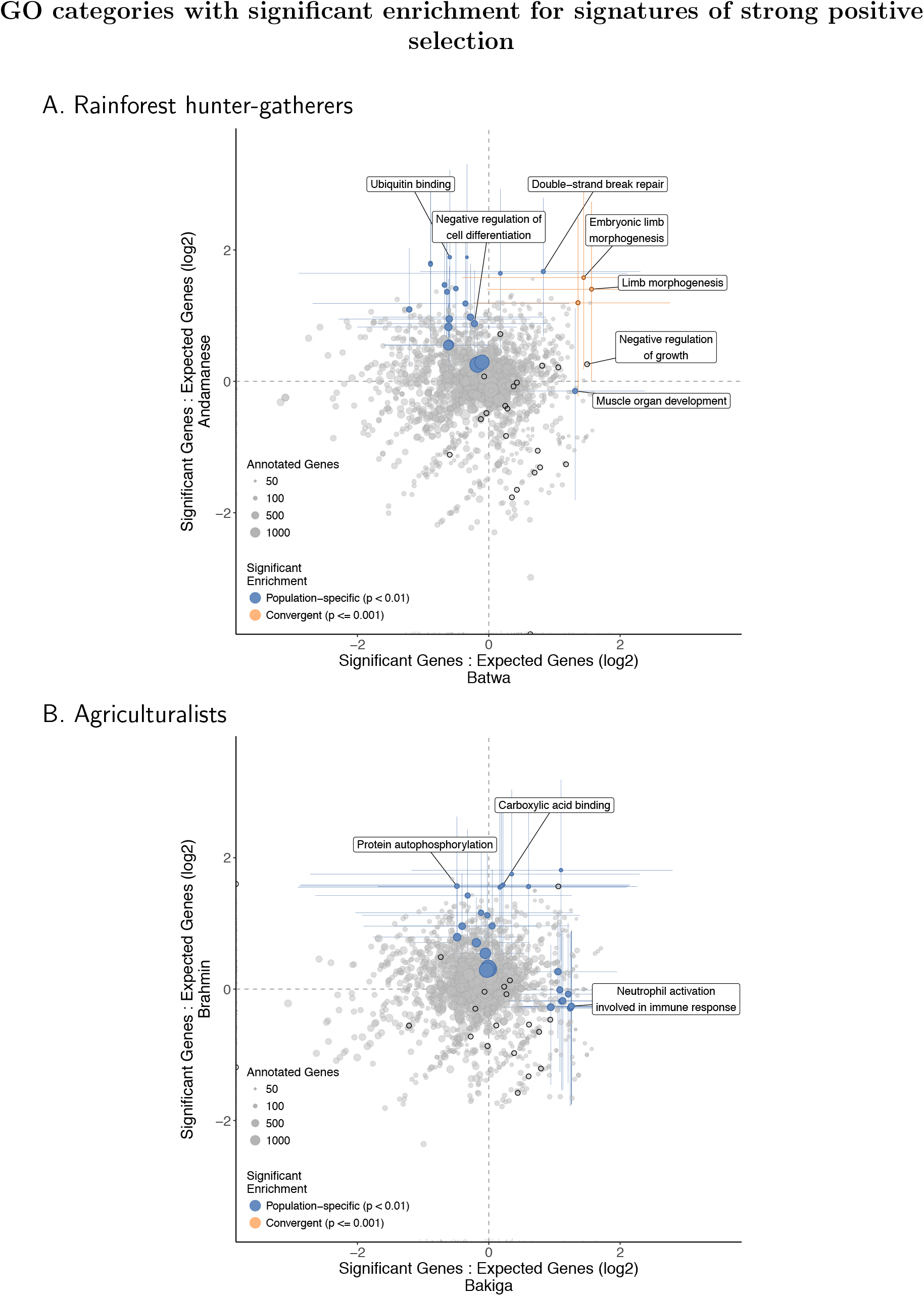
Gene Ontology (GO) functional categories’ ratios of expected to observed counts of outlier genes (with PBS selection index < 0.01) in the Batwa and Andamanese rainforest hunter-gatherers (A) and Bakiga and Brahmin agriculturalist control comparison (B). Results shown for GO biological processes and molecular functions. Point size is scaled to number of annotated genes in category. Terms that are significantly overrepresented for genes under positive selection (Fisher *p* < 0.01) in either population are shown in blue and for both populations convergently (empirical permutation-based *p* ≤ 0.001) are shown in orange. Colored lines represent 95% CI for significant categories estimated by bootstrapping genes within pathways. Dark outlines indicate growth-associated terms: the ‘growth’ biological process (GO:0040007) and its descendant terms, or the molecular functions ‘growth factor binding,’ ‘growth factor receptor binding,’ ‘growth hormone receptor activity,’ and ‘growth factor activity’ and their sub-categories.

Other functional categories of genes were overrepresented in the sets of outlier loci for one of these hunter-gatherer populations but not the other (Fig. 1A; Table S2, S24). The top population-specific enrichments for genes with outlier PBS selection index values for the Batwa were associated with growth and development: ‘muscle organ development’ (GO:0007517; observed genes: 10; expected genes: 4.02; *p* = 0.007) and ‘negative regulation of growth’ (GO:0045926; observed = 7; expected = 2.48; *p* = 0.012). Significantly overrepresented GO biological processes for the Andamanese included ‘negative regulation of cell differentiation’ (GO:0045596; observed genes: 18; expected genes: 9.79; *p* = 0.009). However, these population-specific enrichments were not significant following multiple test correction (false discovery rate *q* = 0.71 for both Batwa terms and *q* = 0.22 for the An-A. Rainforest hunter-gatherers damanese result).

In contrast, no GO functional categories were observed to have similarly significant convergent excesses of ‘outlier’ genes with signatures of positive selection across the two agriculturalist populations as that observed for the rainforest hunter-gatherer populations (Fig. 1B; Table S19), and the top ranked GO categories from both the convergent evolution analysis and the population-specific analyses were absent any obvious connections to skeletal growth. The top-ranked functional categories with enrichments for genes with outlier PBS selection index values for the individual agriculturalist populations included ‘neutrophil activation involved in immune response’ for the Bakiga (GO:0002283; observed = 13; expected = 5.43; *p* = 0.003; *q* = 0.41) and ‘protein autophosphorylation’ for the Brahmin (GO:0046777; observed = 11; expected = 3.71; *p* = 0.0012; *q* = 0.16; Table S24).

### Signatures of convergent and population-specific polygenic adaptation

Outlier-based approaches such as that presented above are expected to have limited power to identify signatures of polygenic adaptation [7–11], which is our expectation for the pygmy phenotype [17]. Unlike the previous analyses in which we identified functional categories with an enriched number of genes with outlier PBS selection index values, for our polygenic evolution analysis we computed a “distribution shift-based” statistic to instead identify functionally-grouped sets of loci with relative shifts in their distributions of PBS selection indices. Specifically, we used the Kolmogorov-Smirnov (KS) test to quantify the distance between the distribution of PBS selection indices for the genes within a functional category to that of the genome-wide distribution. Significantly positive shifts in the PBS selection index distribution for a particular functional category may reflect individually subtle but consistent allele frequency shifts across genes within the category, which could result from either a relaxation of functional constraint or a history of polygenic adaptation. Our approach is similar to another recent method that was used to detect polygenic signatures of pathogen-mediated adaptation in humans [31]. As above, we identified functional categories with convergently high KS values between cross-continental groups by repeating these tests 1,000 times on permuted gene-PBS values and computing the joint probability of both null hypotheses being false for the two populations. We then compared this value from the random iterations to the same statistic computed with the observed KS p-values for each functional category. For example, for the Batwa and Andamanese, we tallied the number of random iterations for which the joint probability of both null hypotheses being false was more extreme (lower) than those of the random iterations. In this way we tested the null hypothesis that both of our observed p-values could have been jointly generated by random chance.

The GO molecular function with the strongest signature of a convergent polygenic shift in PBS selection indices across the Batwa and Andamanese populations was ‘growth factor binding’ (Table S3; Fig. 2A; GO:0019838; Batwa *p* = 0.021; Andamanese *p* = 0.027; Fisher’s combined *p* = 0.0048; empirical test for convergence *p* < 0.001; *q* < 0.001), and the top GO biological process was ‘organ growth’ (GO:0035265; Batwa *p* = 0.028; Andamanese *p* = 0.045; Fisher’s combined *p* = 0.0095; empirical test for convergence *p* = 0.001; *q* = 1). The other top Batwa-Andamanese convergent GO biological processes are not as obviously related to growth, but instead involve muscles, particularly heart muscles. A significant convergent shift in PBS selection indices across both hunter-gatherer populations was observed for ‘cardiac muscle tissue development’ (GO:0048738; Batwa *p* = 0.046; Andamanese *p* = 0.003; Fisher’s combined *p* = 0.001; empirical test for convergence *p* = 0.001; *q* = 1).

**Figure 2:**
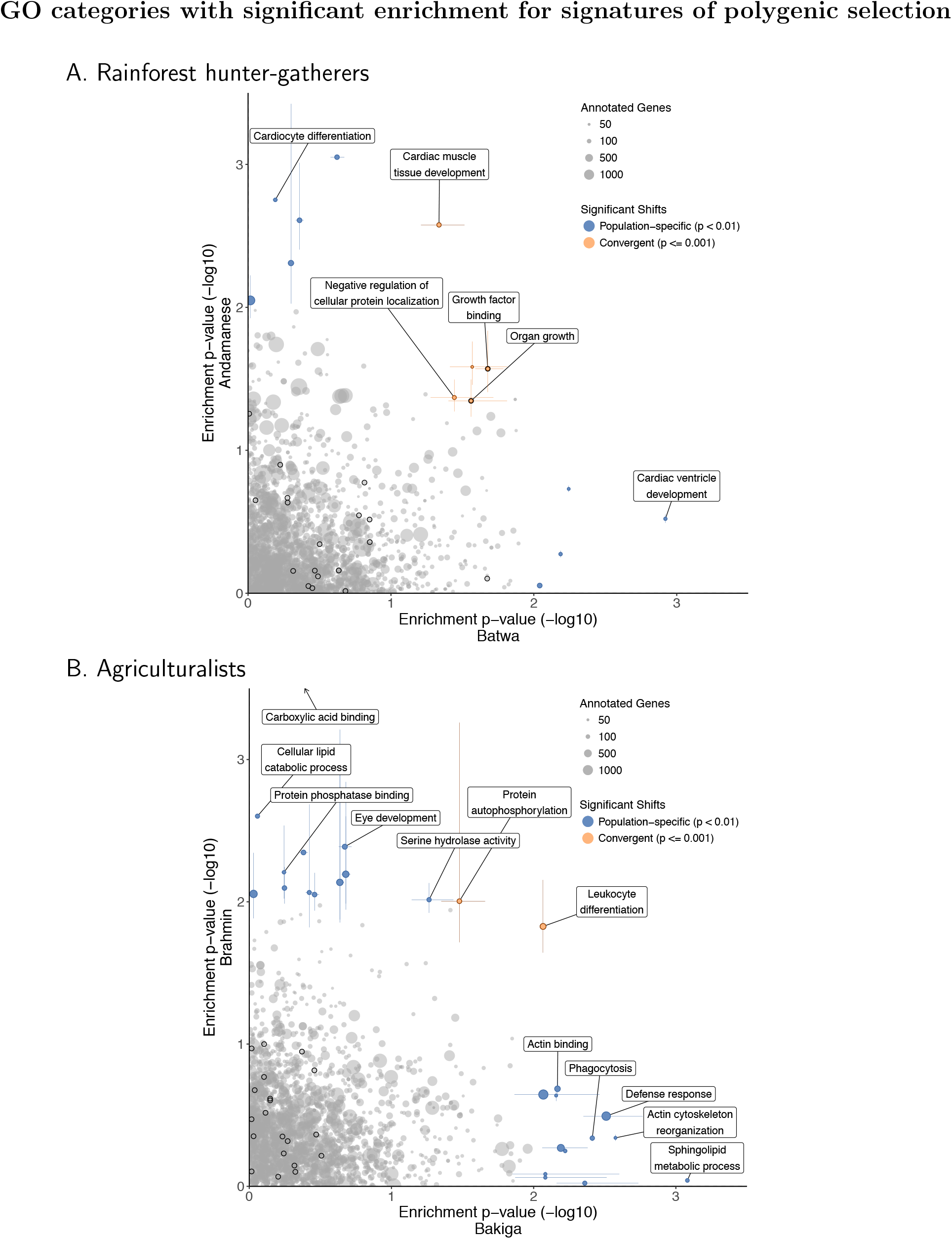
Gene Ontology (GO) functional categories’ distribution shift test p-values, indicating a shift in the PBS selection index values for these genes, in the Batwa and Andamanese rainforest hunter-gatherers (A) and Bakiga and Brahmin agriculturalist control comparison (B). Results shown for GO biological processes and molecular functions. Point size is scaled to number of annotated genes in category. Terms that are significantly enriched for genes under positive selection (Kolmogorov-Smirnov *p* < 0.01) in either population are shown in blue and for both populations convergently (empirical permutation-based *p* ≤ 0.001) are shown in orange. Colored lines represent 95% CI for significant categories estimated by bootstrapping genes within pathways. Dark outlines indicate growth-associated terms: the ‘growth’ biological process (GO:0040007) and its descendant terms, or the molecular functions ‘growth factor binding,’ ‘growth factor receptor binding,’ ‘growth hormone receptor activity,’ and ‘growth factor activity’ and their sub-categories. 0ne GO molecular function, “carboxylic acid binding” (GO:0031406; Brahmin *p* =7.3 × 10^−5^; *q* = 0.0050) not shown, but indicated with arrow.

In contrast, when this analysis was repeated on the agriculturalist populations, no growth-or muscle-related functional annotations were observed with significantly convergent shifts in both populations (Fig. 2B; Table S26). The GO categories with evidence of potential convergent evolution between the agriculturalists were the biological processes ‘leukocyte differentiation’ (GO:0002521; Bakiga *p* = 0.0086; Brahmin *p* = 0.0149; Fisher’s combined *p* = 0.00128; convergence empirical *p* < 0.001; *q* < 0.001) and ‘protein autophosphorylation’ (GO:0046777; Bakiga *p* = 0.033; Brahmin *p* = 0.0099; Fisher’s combined *p* = 0.003; convergence empirical *p* = 0.001; *q* =1).

We also used Bayenv, a Bayesian linear modeling method for identifying loci with allele frequencies that covary with an ecological variable [9, 32], to assess the level of consistency with our convergent polygenic PBS shift results. Specifically, we used Bayenv to test whether the inclusion of a binary variable indicating subsistence strategy would increase the power to explain patterns of genetic diversity for a given functional category of loci over a model that only considered population history (as inferred from the covariance of genome-wide allele frequencies in the dataset.) We converted Bayes factors into per-gene index values via permutation of SNP-gene associations (Table S21) and identified GO terms with significant shifts in the Bayenv Bayes factor index distribution [9, 32] (Table S27). The top results from this analysis included ‘growth factor activity’ (GO:0008083; *p* = 0.006; *q* = 0.11), categories related to enzyme regulation (e.g. ‘enzyme regulator activity’; GO:0030234; *p* = 0.003; *q* = 0.01), and categories related to muscle cell function (e.g. ‘microtubule binding’; GO:0008017; *p* = 0.003; *q* = 0.10). There were more GO terms that were highly ranked (*p* < 0.05) in both the hunter-gatherer PBS shift-based empirical test of convergence and the Bayenv analysis than expected by chance (for biological processes GO terms: observed categories in common = 13, expected = 8.03, Fisher’s exact test *p* = 9.67 × 10^−5^; for molecular function GO terms: observed categories in common = 4, expected = 1.45, Fisher’s exact test *p* = 0.045).

While we did not observe any significant population-specific shifts in PBS selection index values for growth-associated GO functional categories in any of our studied populations (Table S4; Suppl. Text), for each individual rainforest hunter-gatherer population we did observe nominal shifts in separate biological process categories involving the heart (Fig. 2A). For the Batwa, ‘cardiac ventricle development’ (GO:0003231) was the top population-specific result (median PBS index = 0.272 vs. genome-wide median PBS index = 0.528; *p* = 0.001; *q* = 0.302). For the Andamanese, ‘cardiocyte differentiation’ (GO:0035051) was also ranked highly (median PBS index = 0.353 vs. genome-wide median PBS index = 0.552; *p* = 0.002; *q* = 0.232). We note that while these are separate population-specific signatures, 17 genes are shared between the above two cardiac-related pathways (of 61 total ‘cardiocyte differentiation’ genes total, 28%; of 71 total ‘cardiac ventricle development’ genes, 24%; Table S28).

In contrast, cardiac development-related GO categories were not observed among those with highly-ranked population-specific polygenic shifts in selection index values for either the Bakiga or Brahmin agriculturalists (Fig. 2B; Table S29). The only GO term with a significant population-specific shift in the agriculturalists after multiple test correction was molecular function ‘carboxylic acid binding’ in the Brahmins (GO:0031406; *p* = 7.30 × 10^−5^; *q* = 0.005).

To ensure that our results were robust to several possible biases, we repeated the above analyses with several modifications. First, to control for potential biases related to variation in gene length and SNP minor allele frequency (MAF), we repeated all analyses after computing the PBS selection index with binning of genes by length and SNPs by MAF, respectively. Our results were not materially different (Tables S5-S12; Figs. S4-S8; Suppl. Text). Second, to account for the effect of linkage disequilibrium among SNPs within a gene, we re-computed the empirical test for convergence p-values by permuting gene-GO relationships when generating the random null distributions for the PBS selection index values instead of gene-PBS relationships as in our original analysis. Again, downstream results were largely unchanged (Table S13-S14; Suppl. Text). These additional analyses increase our confidence that our results are not artifactual.

## Discussion

The independent evolution of small adult body size in multiple different tropical rainforest environments worldwide presents a natural human model for comparative study of the genetic and evolutionary bases of growth and body size. Through an evolutionary genomic comparison of African and Asian rainforest hunter-gatherer populations to one another and with nearby agriculturalists, we have gained additional, indirect insight into the genetic structure of body size, a fundamental biological trait. Specifically, we identified a signature of potential convergent positive selection on the growth factor binding pathway that could partially underlie the independent evolution of small body size in African and Asian rainforest hunter-gatherers.

Unexpectedly, we also observed signatures of potential polygenic selection across functional categories of genes related to heart development in the rainforest hunter-gatherer populations, both convergently and on a population-specific basis. To a minor extent, the growth factor-and heart-related functional categories highlighted in our study do overlap: of the 123 total genes annotated across the three heart-related categories (‘cardiac muscle tissue development’ GO:0048738, ‘cardiac ventricle development’ GO:0003231, and ‘cardiocyte differentiation’ GO:0035051), nine (7.3%) are also included among the 66 annotated genes in the ‘growth factor binding’ category (GO:0019838). However, even after excluding these nine genes from our dataset, we still observed similar polygenic PBS shifts in the Batwa and Andamanese for both growth factor-and heart-related functional categories (Suppl. Text), demonstrating that our observations are not driven solely by cross-annotated genes.

We hypothesize that the evolution of growth hormone sub-responsiveness, which appears to at least partly underlie short stature in some rainforest hunter-gatherer populations [33–37] may in turn have also resulted in strong selection pressure for compensatory adaptations in cardiac pathways. The important roles of growth hormone (GH1) in the heart are evident from studies of patients deficient in the hormone. For example, patients with growth hormone deficiency are known to be at an increased risk of atherosclerosis and mortality from cardiovascular disease [38] and have worse cardiac function [39]. More broadly, shorter people have elevated risk of coronary artery disease [40], likely due to the pleiotropic effects of variants affecting height and atherosclerosis development [41]. Such health outcomes may relate to the important roles that growth hormone plays in the development and function in the myocardium [42, 43], which contains a relatively high concentration of receptors for growth hormone [44]. We hypothesize that the adaptive evolution of growth hormone subresponsiveness underlying short stature in rainforest hunter-gatherers may have necessitated compensatory adaptations in the cardiac pathways reliant on growth hormone.

An alternative explanation for our finding of potential convergent positive selection on cardiac-related pathways relates to the nutritional stress of full-time human rainforest habitation. Especially prior to the ability to trade forest products for cultivated goods with agriculturalists, the diets of full-time rainforest hunter-gatherers may have been calorically and nutritionally restricted on at least a seasonal basis [13]. Caloric restriction has a direct functional impact on cardiac metabolism and function, with modest fasting in mice leading to the depletion of myocardial phospholipids, which potentially act as a metabolic reserve to ensure energy to essential heart functions [45]. In human rainforest hunter-gatherers, selection may have favored variants conferring cardiac phenotypes optimized to maintain myocardial homeostasis during the nutritional stress that these populations may have experienced in the past.

An important caveat to our study is the lack of statistical significance for our population-specific analyses after controlling for the multiplicity of tests resulting from hierarchically nested GO terms. The absence of strong signals of positive selection that are robust to the multiple testing burden likely reflects both the expected subtlety of evolutionary signals of selection on polygenic traits and the restriction of our dataset to gene coding region sequences. However, our comparative approach to identify signatures of convergent evolution is more robust. Therefore, while we cannot yet accurately estimate the extent to which signatures of positive selection that potentially underlie the evolution of the pygmy phenotype occurred in the same versus distinct genetic pathways between the Batwa and Andamanese, we do feel confident in our findings of convergent growth-related and cardiac-related pathways evolution. The concurrent signatures of convergent evolution across these two pathways in both African and Asian rainforest hunter-gatherers is an example of the insight into a biomedically-relevant phenotype that can be gained from the comparative study of human populations with non-pathological natural variation.

## Materials and Methods

### Sample collection and dataset generation

Sample collection, processing, and sequencing have been previously described [17, 23]. Briefly, sampling of biomaterials (blood or saliva) from Batwa rainforest hunter-gatherers and Bakiga agriculturalists of southwestern Uganda took place in 2010 [17]. The study was approved by the Institutional Review Boards (IRBs) of both the University of Chicago (#16986A) and Makerere University, Kampala, Uganda (#2009-137), and local community approval and individual informed consent were obtained before collection. DNA samples of 50 Batwa and 50 Bakiga adults were included in the present study. Exome capture, sequencing, and variant calling were described previously [23]. Briefly, sequence reads were aligned to the hg19/GRCh37 genome with BWA v.0.7.7 mem with default settings [46], PCR duplicates were detected with Picard Tools v.1.94 (http://broadinstitute.github.io/picard), and re-alignment around indels and base quality recalibration was done with GATK v3.5 [47] using the known indel sites from the 1000 Genomes Project [26]. Variants were called individually with GATK HaplotypeCaller [47], and variants were pooled together with GATK GenotypeGVCF and filtered using VQSR. Only biallelic SNPs with a minimum depth of 5x and less than 85% missingness that were polymorphic in the entire dataset were retained for analyses.

Variant data for the Andamanese individuals (Jarawa and Onge) and an outgroup mainland Indian population (Uttar Pradesh Brahmins) from [25] were downloaded in VCF file format from a public website. To ensure the exome capture-derived African and whole genome shotgun sequencing-derived Asian datasets were comparable, we restricted our analyses of these data to exonic SNPs only.

### Merging with 1000 Genomes data

We chose outgroup comparison populations from the 1000 Genomes Project [26] to be equally distantly related to the ingroup populations: Reads from a random sample of 30 unrelated individuals from British in England and Scotland (GBR) and Luhya in Webuye, Kenya (LWK) were chosen for the Batwa/Bakiga and Andamanese/Brahmin datasets, respectively. We re-called variants in each 1000 Genomes comparison population at loci that were variable in the ingroup populations using GATK UnifiedGenotyper [47]. Variants were filtered to exclude those with QD < 2.0, MQ < 40.0, FS > 60.0, HaplotypeScore > 13.0, MQRankSum < −12.5, or ReadPosRankSum < −8.0. We removed SNPs for which fewer than 10 of the 30 individuals from the 1000 Genomes datasets had genotypes.

### Computation of the Population Branch Statistic (PBS) and the per-gene PBS index

Using these merged datasets, we computed *F*_*ST*_ between population pairs using the unbiased estimator of Weir and Cockerham [48], transformed it to a measure of population divergence [*T* = −*1og*(1 − *F*_*ST*_)], and then calculated the Population Branch Statistic (PBS), after [29]. PBS was computed on a per-SNP basis. We computed an empirical p-value for each SNP, simply the proportion of coding SNPs with PBS greater than the value for this SNP, which we adjusted for FDR.

SNPs were annotated with gene-based information using ANNOVAR [49] with refGene (Release 76) [50] and PolyPhen [51] data. As the Andamanese/Brahmin dataset spanned the genome and the Batwa/Bakiga exome dataset included off target intronic sequences as well as untranslated regions (UTRs), and microRNAs, we restricted our analysis to only exonic SNPs. For both the Batwa/Bakiga and Andamanese/Brahmin datasets, we computed a “PBS selection index” for each gene as follows. We compared the mean PBS for all SNPs located within that gene to a distribution of values estimated by shuffling SNP-gene associations (without replacement) and re-computing the mean PBS value for that gene
10,0 times. We defined the PBS selection index of the gene as the percentage of these empirical mean values that is higher than its observed mean PBS value. When identifying outlier genes, gene-based indices were adjusted for FDR.

In order to assess potential biases related to variation in gene length and SNP minor allele frequencies (MAF), we repeated all analyses after computing the PBS selection index with binning of genes by length or SNPs by MAF. Complete details of these methods are included in the Supplemental Text.

To identify SNPs with allele frequencies correlated with subsistence strategy (hunter-gatherer: Andamanese and Batwa; agriculturalists: Bakiga and Brahmin), we used Bayenv2.0 [32] to assess whether the addition of a binary variable denoting subsistence strategy improved the Bayesian model that already took into account covariance between samples due to ancestry. As with the PBS results, we computed an index for each gene by sampling new values for each SNP from the distribution of all Bayes factors and comparing the actual average for this gene to those of the bootstrapped replicates.

### Creation of *a priori* lists of growth-related genes

To test the hypothesis that genes with known influence on growth would show increased positive selection in rainforest hunter-gatherer populations, we curated *a priori* lists of growth-related genes as described fully in the Supplemental Text. Briefly, we obtained the following gene lists: i) 3,996 genes that affect growth or size in mice (MP:0005378) from the Mouse/Human Orthology with Phenotype Annotations database [52]; ii) 266 genes associated with abnormal skeletal growth syndromes in the Online Mendelian Inheritance in Man (OMIM) database (https://omim.org), as assembled by [53]; iii) 427 genes expressed substantially more highly in the mouse growth plate, the cartilaginous region on the end of long bones where bone elongation occurs, than in soft tissues [lung, kidney, heart; >= 2.0 fold change; [54]]; and iv) 955 genes annotated with the Gene Ontology “growth” biological process (GO:0040007). As the GH/IGF1 pathway is a major regulator of growth and disruptions to the pathway have been implicated in the pygmy phenotype, we also collected lists of genes associated with GH1 and IGF1 respectively from the OPHID database of pro-teinprotein interaction (PPI) networks [55]. Separately, we also used a list of genes found to be associated with the pygmy phenotype in the Batwa [17].

### Statistical overrepresentation and distribution shift tests

Using the PBS and Bayenv indices, we next tested for a statistical over-representation of extreme values (*p* < 0.01) for the above *a priori* gene lists as well as all Gene Ontology (GO) terms using the topGO package of Bioconductor [56]s, gene-to-GO mapping from the org.Hs.eg.db package [57], and Fisher’s exact test in “classic” mode (i.e., without adjustment for GO hierarchy). We similarly performed a statistical enrichment test using the Kolmogorov-Smirnov test again in “classic” mode, which tested for a shift in the distribution of the PBS or Bayenv statistic, rather than an excess of extreme values. In all cases, we pruned the GO hierarchy to exclude GO terms with fewer than 50 annotated genes to reduce the number of tests, leaving 1,742 and 1,816 GO biological processes and 266 and 285 GO molecular functions tested for the African and Asian datasets, respectively. To further reduce the number of redundant tests, we also computed the semantic similarity between GO terms to remove very similar terms. We computed the similarity metric of [58] as implemented in the GoSemSim R package [59] a measure of the overlapping information content in each term using the annotation statistics of their common ancestor terms, and then clustered based on these pairwise distances between GO terms using Ward Hierarchical Clustering. We then pruned GO terms by cutting the tree at a height of 0.5 and retaining the term in each cluster with the lowest p-value. With this reduced set of GO overrepresentation and distribution shift results, we adjusted the p-value for FDR.

### Identification of signatures of convergent evolution

We used two methods to identify convergent evolution: i.) computation of simple combined p-values for SNPs, genes, and GO overrepresentation and distribution shift tests using Fisher’s and Edgington’s methods, and ii.) a permutation based approach to identify GO pathways for which both the Batwa and Andamanese overrepresentation or distribution shift test results are more extreme than is to be expected by chance (the “empirical test for convergence”). These two approaches are summarized below.

We searched for convergence between Batwa and Andamanese individuals by computing the joint p-value for PBS on a per-SNP, per-gene, and per-GO term basis. We calculated all joint p-values using Fisher’s method (as the sum of the natural logarithms of the uncorrected p-values for the Batwa and Andamanese tests [60]) as well as via Edgington’s method (based on the sum of all p-values [61]). Meta-analysis of p-values was done via custom script and the metap R package [62].

We also assessed the probability of getting two false positives in the Batwa and Andamanese selection results by shuffling the genes’ PBS indices 1,000 times and performing GO overrepresentation and distribution shift tests on these permuted values. We compared the observed Batwa and Andamanese p-values to this generated distribution of p-values, as described above. We computed the joint probability of both null hypotheses being false for the Andamanese and Batwa as (1 − *p*_*Batwa*_)(1− *p*_*Andamanese*_), where *p*_*Batwa*_ and *p*_*Andamanese*_ are the p-values of the Fisher’s exact test or of the Kolmogorov-Smirnov test for the outlier-and shift-based tests, respectively, and we compared the joint probability to the same statistic computed for the p-values from the random iterations. The empirical test for convergence p-value was simply the number of iterations for which this statistic was more extreme (lower) for the observed values than for the randomly generated values.

We also performed a variation of this analysis, but to preserve patterns of linkage disequilibrium among SNPs within a gene in the null distribution, instead of permuting gene-PBS relationships to generate the random null distributions for the PBS selection index values of the two populations considered jointly, we instead permuted the gene-GO relationships. That is, to compute the PBS selection index, the one-to-many relationships between genes and GO terms were shuffled when generating the null distribution, maintaining the groupings of GO terms that were assigned together to an original gene. Full details of this analysis are available in the Supplemental Text.

### Script and data availability

All scripts used in the analysis are available at https://github.com/bergeycm/rhg-convergence- analysis and released under the GNU General Public License v3. Exome data for the Batwa and Bakiga populations have previously been deposited in the European Genome-phenome Archive under accession code EGAS00001002457. Extended data tables are available at https://doi.org/10.18113/S1N63M.

## Acknowledgments

The authors would like to thank the Batwa and Bakiga communities and all individuals who participated in this study, and J.A. Hodgson and E.C. Reeves for helpful discussions. This work was supported by NIH R01-GM115656 (to G.H.P and L.B.B.), 1 F32 GM125228-01A1 (to C.M.B), and ANR AGRHUM ANR-14-CE02-0003-01 (to L.Q.-M.). M.L. was supported by the Fondation pour la Recherche Médicale (FDT20170436932). This research was conducted with Advanced CyberInfrastructure computational resources provided by The Institute for CyberScience at The Pennsylvania State University.

## 1 Supplemental Text

### Positive selection signatures on growth-associated genes

We examined whether gene-specific signatures of strong positive selection (using an “outlier-based”” designation of genes with PBS index values < 0.01) in the rainforest populations were enriched for known functional associations with growth using *a priori* lists of 4,888 total growth-related genes, consisting of (with some redundancy among individual categories, as expected): i) 3,996 genes that affect growth or size in mice (MP:0005378) from the Mouse/Human Orthology with Phenotype Annotations database [1]; ii) 266 genes associated with abnormal skeletal growth syndromes in the Online Mendelian Inheritance in Man (OMIM) database (https://omim.org/), as assembled by [2]; iii) 427 genes expressed substantially more highly in the mouse growth plate, the cartilaginous region on the end of long bones where bone elongation occurs, than in soft tissues (lung, kidney, heart; >= 2.0 fold change; [3]; and iv) 955 genes annotated with the Gene Ontology “growth” biological process (GO:0040007). Separately, we also considered in our analyses the set of 166 genes located within the 16 genomic regions previously associated with the pygmy phenotype in the Batwa, using an admixture mapping approach [4], as well as GH1-and IGF1-associated genes using data from OPHID database of proteinprotein interaction (PPI) networks [5].

We used each of the curated *a priori* growth-related gene lists for testing the hypothesis that such loci are enriched for genes with signatures of strong positive selection (outlier PBS selection index values) or have a shift in the distribution of PBS selection index values consistent with subtle polygenic adaptation in the Batwa and Andamanese rainforest hunter-gatherer but not the Bakiga and Brahmin agriculturalist populations. We identified 202, 188, 291, and 252 outlier strong selection candidate genes (with PBS index values < 0.01) in each of the Batwa, Bakiga, Andamanese, and Brahmin populations, respectively. Genes in the *a priori* growth-related gene lists were not significantly overrepresented among PBS outliers in any populations, except for those associated with mouse growth phenotype in the Brahmin (68 observed, 47.7 expected; Fisher p = 0.0179) (Table S23). Though the lack of over-representation of growth-related gene lists among loci with outlier signatures of strong positive natural selection related to growth is perhaps unsurprising considering the polygenic phenotype, our distribution shift-based test also showed no significant shifts in the distribution of PBS indices for any population (Table S25). Genes in genomic regions previously associated with the pygmy phenotype in the Batwa [4] were enriched for genes with outlier PBS selection index values in the Batwa (outlier-based test: 5 observed, 1.39 expected; Fisher *p* = 0.017; Table S23) and the PBS distribution for the phenotype-associated genes was shifted relative to the genome-wide distribution (distribution shift-based test: Kolmogorov-Smirnov test *p* = 0.056; Table S25). We found no evidence that genes associated with GH1 and IGF1 were enriched for outlier or polygenic selection.

### Impact of cross-annotated genes between growth factor- and cardiac-related pathways

To assess whether shared genes in GO categories relating to the heart and growth factor binding were responsible for the significant shift in PBS selection index values for genes in these annotations, we compared the distributions of PBS selection indices before and after removing 9 genes common to heart pathways and growth factor binding. The heart GO terms assessed were: ‘cardiocyte differentiation’ (GO:0035051), with a shift in the Andamanese hunter-gatherers; ‘cardiac ventricle development’ (GO:0003231), with a shift in the Batwa hunter-gatherers; and ‘cardiac muscle tissue development’ (GO:0048738) with a convergent shift in the Batwa and Andamanese. Of the 123 heart related genes contained in these pathways, 9 were also annotated to the GO molecular function ‘growth factor binding’ (GO:0019838): *ACVR1, EGFR, ENG, FGFR2, FGFRL1, LTBP1, SCN5A, TGFBR1*, and *TGFBR3.*

After removing the 9 shared genes, the mean PBS selection index for the Andamanese among genes annotated to ‘cardiocyte differentiation’ decreased slightly from 0.444 to 0.443 and the pre-and post-filtration distributions were not significantly different (Kolmogorov-Smirnov *D* = 0.023, *p* =1). Similarly, the mean PBS selection index for the Batwa for genes in ‘cardiac ventricle development’ decreased slightly from 0.654 to 0.652, and the distributions were not significantly different (*D* = 0.044, *p* = 1). Finally, for ‘cardiac muscle tissue development’, the mean PBS selection index for the Andamanese increased from 0.450 to 0.453, and for the Batwa increased from 0.474 to 0.486. Again the pre-and post-filtering distributions were not significantly different for the Andamanese (*D* = 0.015, *p* = 1) or Batwa (*D* = 0.015, *p* = 1).

Similarly, after removing 9 shared genes, the mean PBS selection index for genes annotated to ‘growth factor binding’ (GO:0019838) for the Batwa increased slightly from 0.437 to 0.440 and for the Andamanese decreased from 0.455 to 0.437. Again, the pairs of distributions were not significantly different (Batwa: *D* = 0.030, *p* =1; Andamanese: *D* = 0.036, *p* = 1).

### Correcting for potential bias from differing gene size or global minor allele frequency (MAF)

In order to assess the potential biases related to differences in gene length (e.g. number of SNPs) or in SNP global minor allele frequencies (MAF), we repeated the analysis after modifying how the PBS selection index was computed. As in the uncorrected analysis, these corrected PBS selection index values were computed using 1,000 iterations.

First, to control for gene size, we sampled the PBS values for each SNP from only genes with the same number of SNPs during the computation of the selection index. For larger genes, gene sizes were binned to ensure sufficient SNPs from which to sample, using sets [11,15], [16, 20], and [21, ∞).

Second, to control for differing MAF values for SNPs, we did the permutation-based computation of the PBS selection index while matching SNPs on global MAF (computed using the African or Asian datasets for within-continent analyses.) SNPs were grouped by MAF into bins of size 0.01, and for each SNP in a gene, SNPs were sampled from only the set in the MAF bin.

Neither modification to the PBS selection index computation algorithm majorly affected the PBS selection index values nor the GO-based downstream analyses. Corrected and uncorrected PBS selection index values were highly correlated (*R*^2^ = 0.993 to 0.997 and 0.953 to 0.985 for the gene size-and MAF-corrections respectively; Fig. S4).

The GO biological processes and molecular functions with the strongest evidence of enrichment for strong selection were similar for the convergent (Tables S5 and S6; Figs. S5 and S6) and population-specific selection analyses (Tables S7 and S8; Figs. S5 and S6). The only mentioned growth-or heart-associated pathway that was no longer significant after correction was the biological process “negative regulation of growth,” which was significantly enriched for genes with evidence of strong selection in the Batwa in the original analysis, but its p-value rose to 0.0448 after correction for gene size. In contrast, “cardiac muscle tissue development” (GO:0048738) which originally had a convergent empirical p-value of 0.025, was significantly enriched for strong positive selection convergently in the Batwa and Andamanese after MAF-based filtration (*p* = 0.001).

Similarly, the top GO categories with evidence of polygenic selection were largely unchanged for the convergent (Tables S9 and S10; Figs. S7 and S8) and population-specific selection analyses (Tables S11 and S12; Figs. S7 and S8). Minor changes include “growth factor binding” (GO:0019838) which rose to be no longer significant with the MAF-based correction (original convergent empirical *p* < 0.001; MAF corrected *p* = 0.005).

### Modification of significance testing in empirical test for convergent evolution

We also modified and repeated the analysis that computes the significance of the convergence GO tests using a permutation-based approach. Whereas we originally permuted gene-PBS relationships to generate the random null distributions of PBS selection index values for two populations considered jointly, we instead permuted the gene-GO relationships to preserve LD patterns. The one-to-many relationships between genes and GO terms were shuffled, maintaining the groupings of GO terms that were assigned together to an original gene. We repeated the GO-based analyses for enrichment of strong selection or polygenic selection times with these randomized gene-GO annotations, and compared our actual observed values to this randomly-generated null distribution. As before, we then defined the p-value of our empirical test for convergent evolution as the probability that this statistic was more extreme (lower) for the observed values than for the randomly generated values. The resultant p-value summarizes the test of the null hypothesis that both results could have been jointly generated under random chance. The results of the modified test were only slightly different than the original for both convergence in strong outlier selection (Table S13) and in a shifted PBS selection index (Table S14).

## 2 Figures

**Fig. S1:**
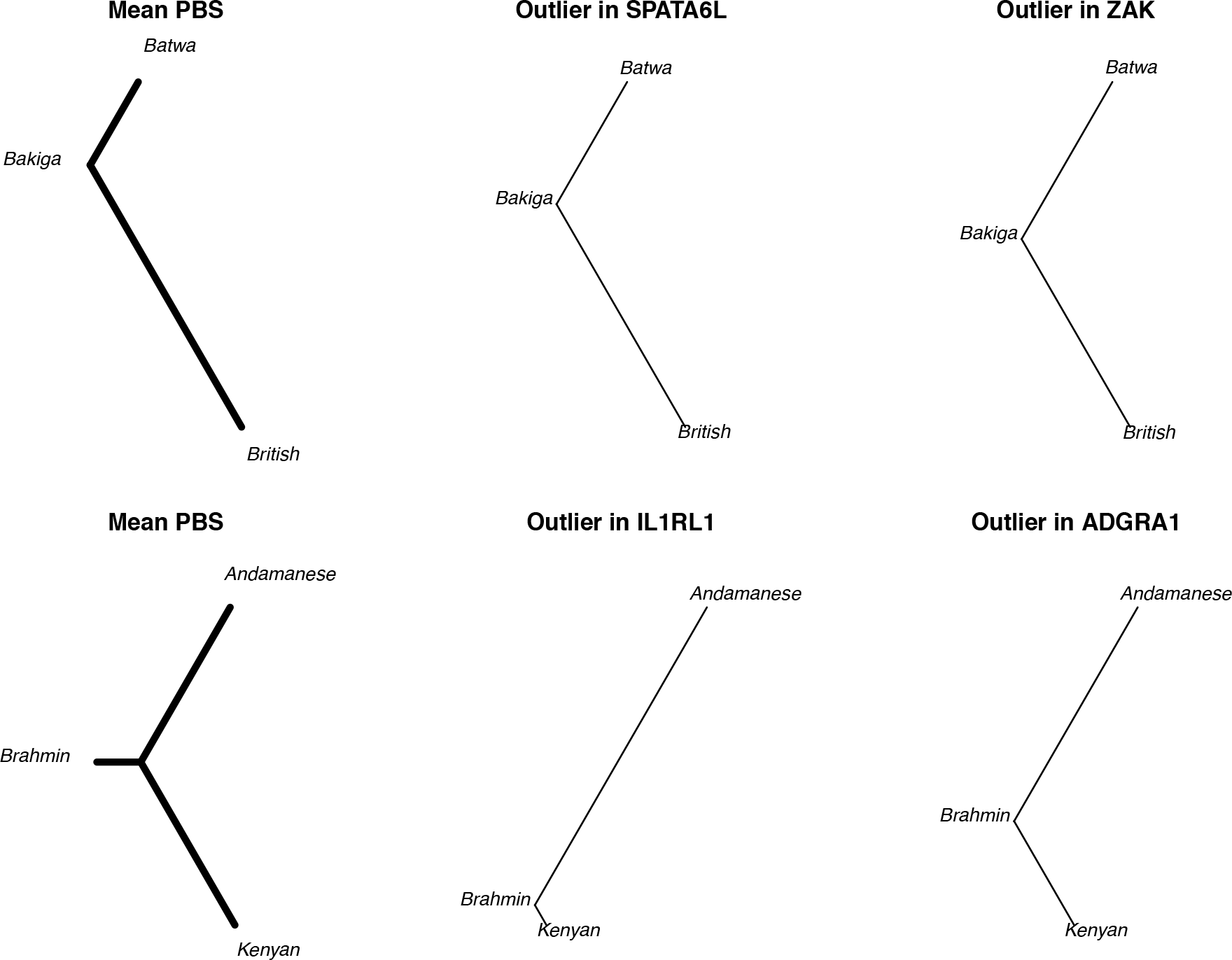
Population Branch Statistic (PBS) schematic. Mean values of the Population Branch Statistic (PBS; left) for the African dataset (Batwa, Bakiga, and outgroup British populations; upper row) and Asian dataset (Andamanese, Brahmin, and outgroup Kenyan populations; lower row). Middle and right columns contain PBS values for two outlier SNPs in each population. Figure S2: Population Branch Statistic (PBS) values plotted across the genome for the four focal populations. The containing the SNPs with the 5 highest PBS values in each population are labeled. Figure S3: Population Branch Statistic (PBS) selection index values plotted by number of SNPs in gene. Color indicates number of genes with that SNP count. Only SNP counts from 1 to 30 shown.

**Fig. S2:**
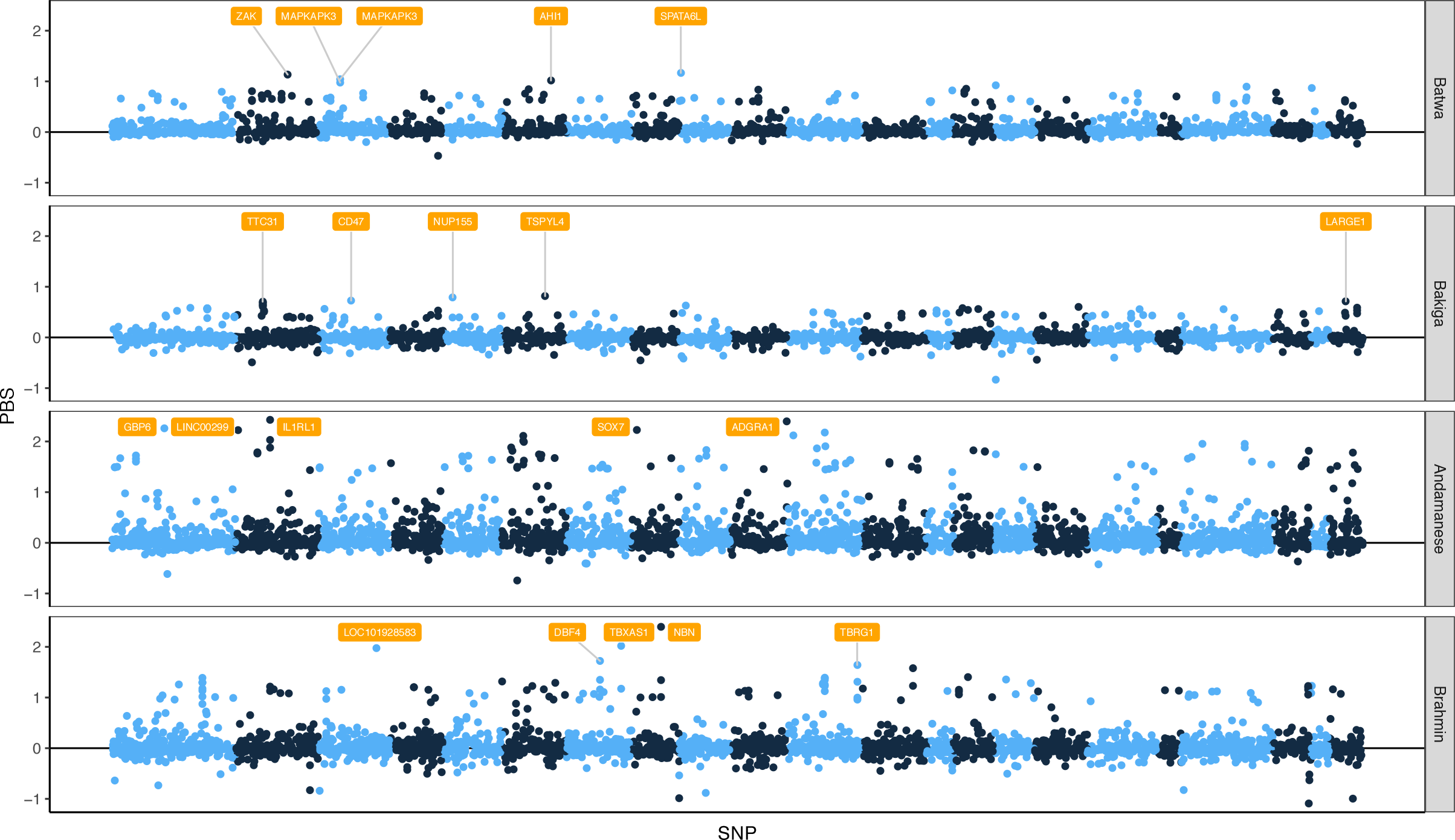
Population Branch Statistic (PBS) by SNP. Population Branch Statistic (PBS) values plotted across the genome for the four focal populations. The genes containing the SNPs with the 5 highest PBS values in each population are labeled.

**Fig. S3:**
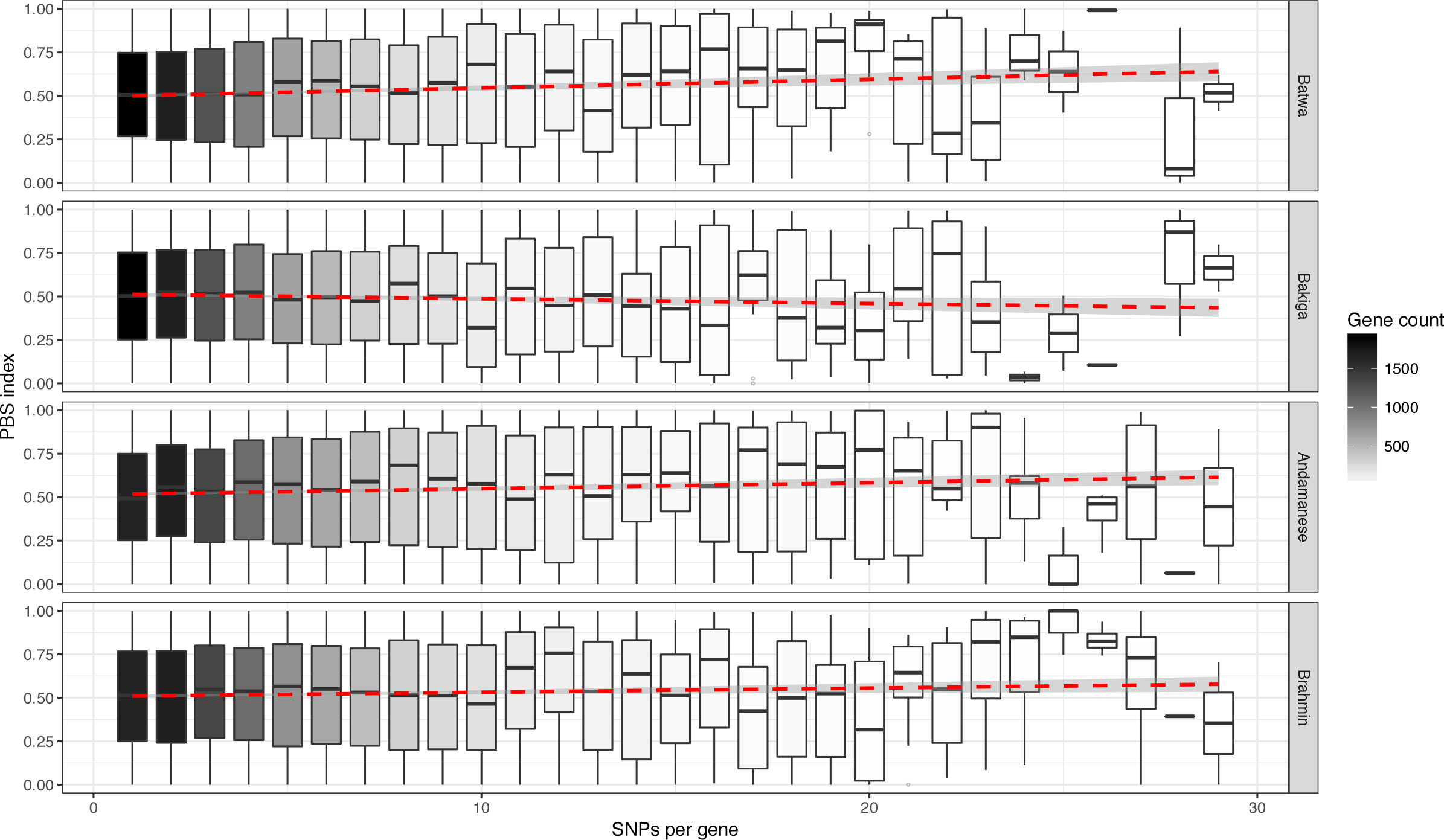
Population Branch Statistic (PBS) by gene SNP count. Population Branch Statistic (PBS) selection index values plotted by number of SNPs in gene. Color indicates number of genes with that SNP count. Only SNP counts from 1 to 30 shown.

**Fig. S4:**
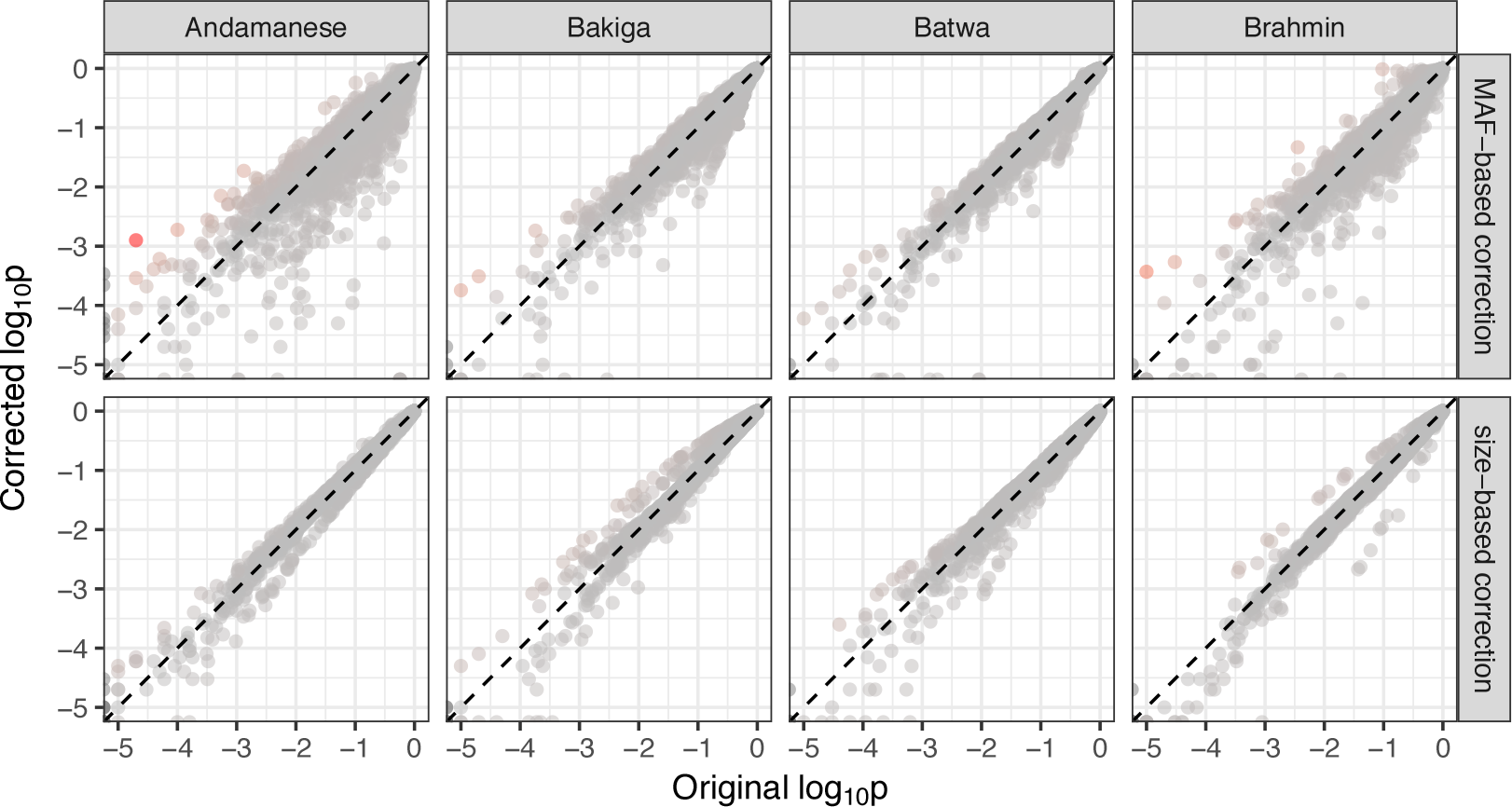
Gene size-and MAF-based corrections impact on p-value. Plots of PBS selection index values for genes corrected for gene size and MAF shown compared to the original uncorrected values (with both plotted on a logarithmic scale. Red shading indicates higher percent difference from originalA. Rainforest hunter-gatherers

**Fig. S5:**
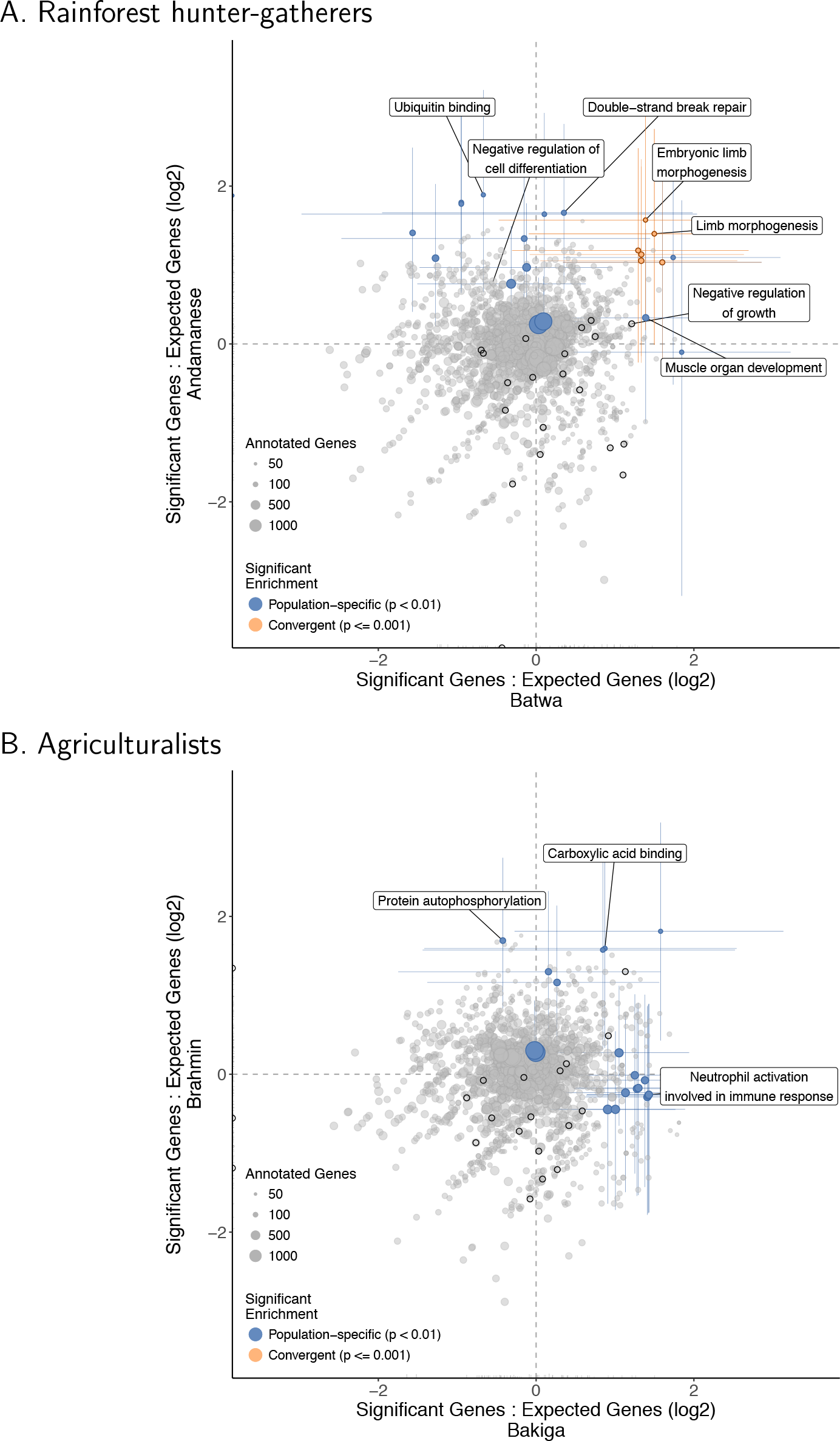
Gene size-corrected strong positive selection enrichment results. After gene size-based correction, Gene Ontology (GO) functional categories’ ratios of expected to observed counts of outlier genes (with PBS selection index < 0.01) in the Batwa and Andamanese rainforest hunter-gatherers (A) and Bakiga and Brahmin agriculturalist control (B). Results shown for GO biological processes and molecular functions. Point size is scaled to number of annotated genes in category. Terms that are significantly overrepresented for genes under positive selection (Fisher *p* < 0.01) in either population shown in blue and for both populations convergently (empirical permutation-based *p* < 0.005) shown in orange. Colored lines represent 95% CI for significant categories estimated by bootstrapping genes within pathways. Dark outlines indicate growth-associated terms: the ‘growth’ biological process (GO:0040007) and its descendant terms, or the molecular functions ‘growth factor binding,’ ‘growth factor receptor binding,’ ‘growth hormone receptor activity,’ and ‘growth factor activity’ and their sub-categories.

**Fig. S6:**
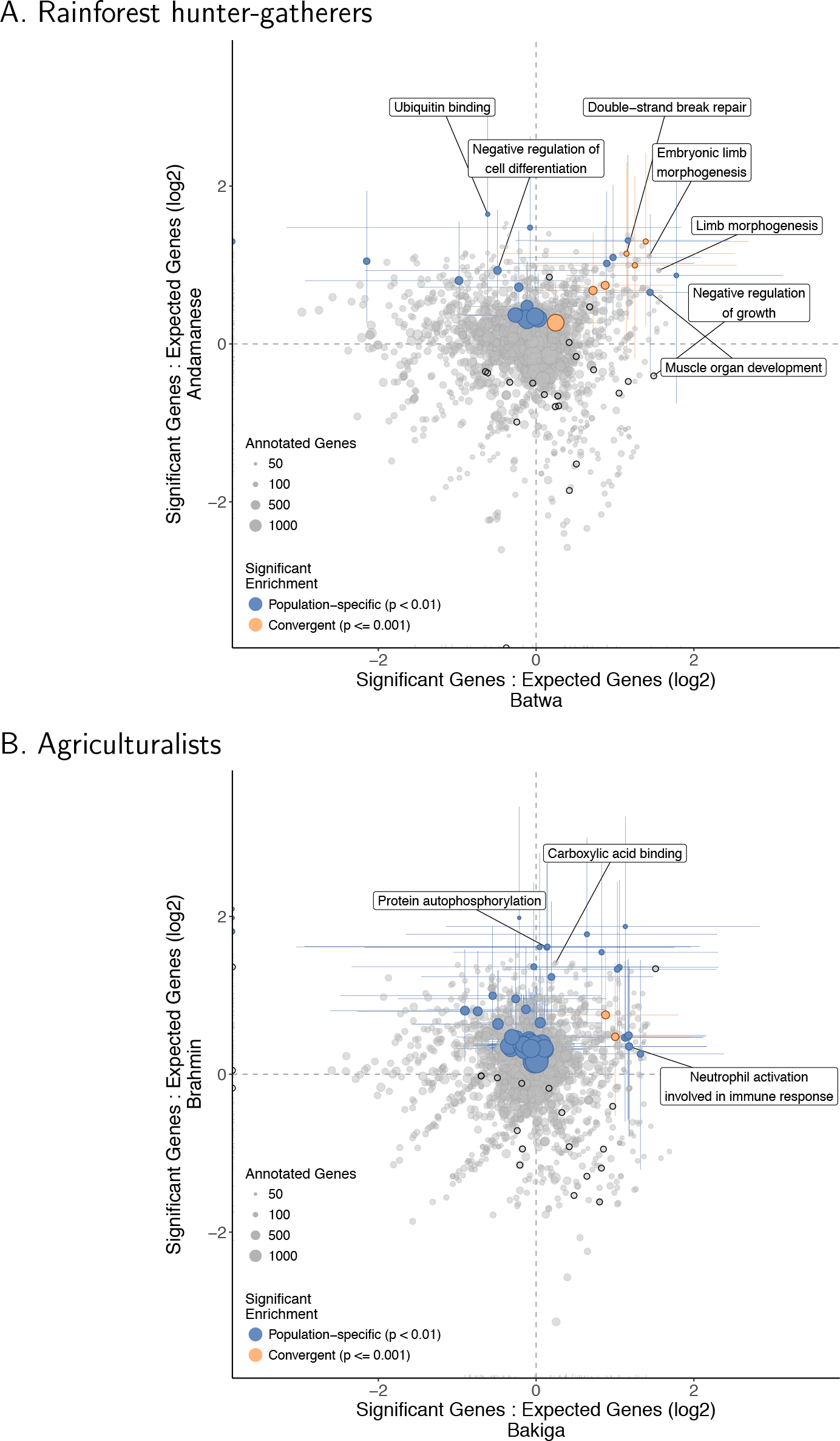
MAF-corrected strong positive selection enrichment results. After MAF-based correction, Gene Ontology (GO) functional categories’ ratios of expected to observed counts of outlier genes (with PBS selection index < 0.01) in the Batwa and Andamanese rainforest hunter-gatherers (A) and Bakiga and Brahmin agriculturalist control (B). Results shown for GO biological processes and molecular functions. Point size is scaled to number of annotated genes in category. Terms that are significantly overrepresented for genes under positive selection (Fisher *p* < 0.01) in either population shown in blue and for both populations convergently (empirical permutation-based *p* < 0.005) shown in orange. Colored lines represent 95% CI for significant categories estimated by bootstrapping genes within pathways. Dark outlines indicate growth-associated terms: the ‘growth’ biological process (GO:0040007) and its descendant terms, or the molecular functions ‘growth factor binding,’ ‘growth factor receptor binding,’ ‘growth hormone receptor activity,’ and ‘growth factor activity’ and their sub-categories.

**Fig. S7:**
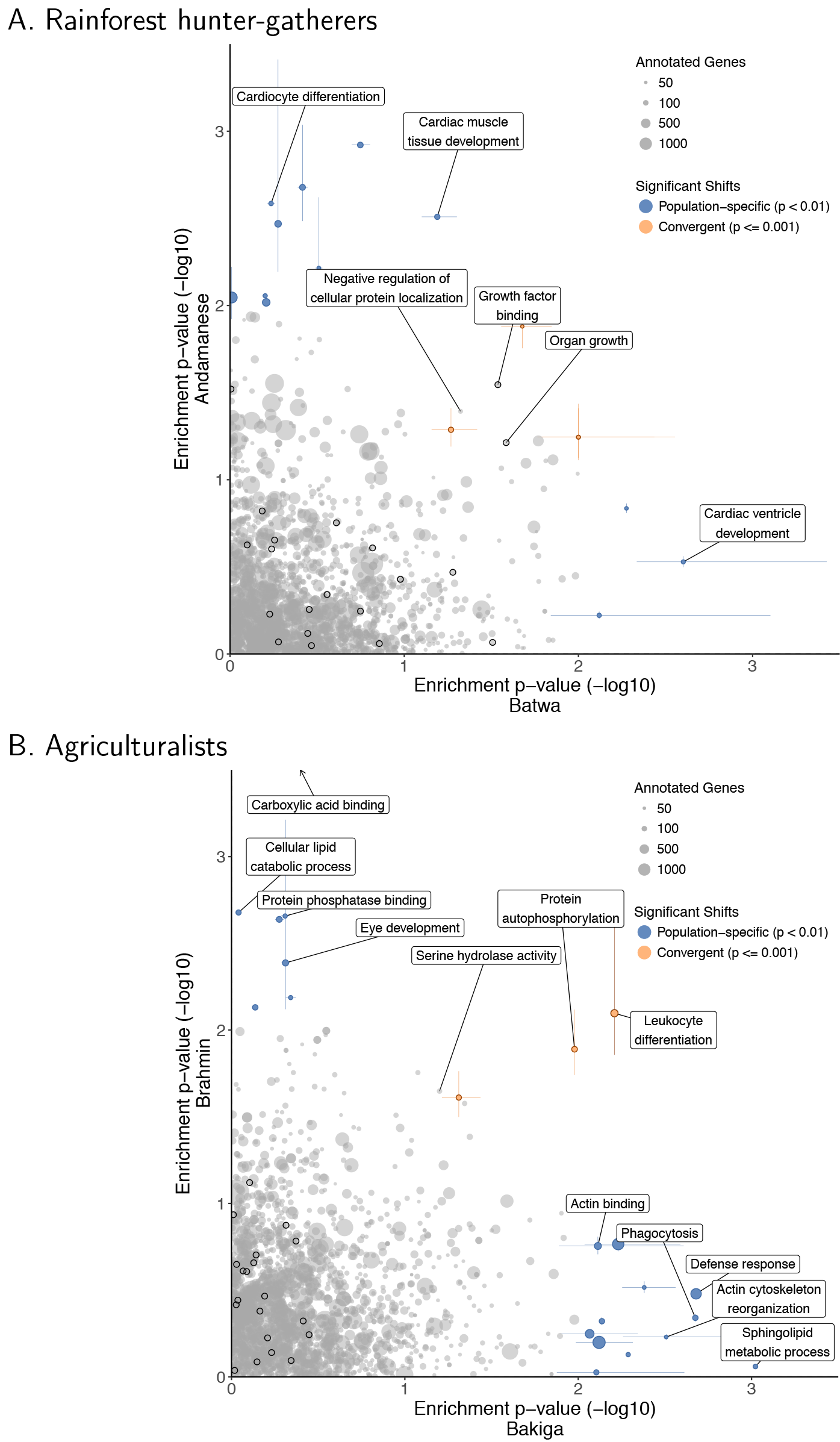
Gene size-corrected polygenic distribution shift test results. After gene size-based correction, Gene Ontology (GO) functional categories’ distribution shift test p-values, indicating a shift in the PBS selection index values for genes, in the Batwa and Andamanese rainforest hunter-gatherers (A) and Bakiga and Brahmin agriculturalist control (B). Results shown for GO biological processes and molecular functions. Point size is scaled to number of annotated genes in category. Terms that are significantly enriched for genes under positive selection (Kolmogorov-Smirnov p < 0.01) in either population shown in blue and for both populations convergently (empirical permutation-based *p* < 0.005) shown in orange. Colored lines represent 95% CI for significant categories estimated by bootstrapping genes within pathways. Dark outlines indicate growth-associated terms: the ‘growth’ biological process (GO:0040007) and its descendant terms, or the molecular functions ‘growth factor binding,’ ‘growth factor receptor binding,’ ‘growth hormone receptor activity,’ and ‘growth factor activity’ and their sub-categories. One GO molecular function, “carboxylic acid binding” (GO:0031406; Brahmin *p* =7.3 × 10^−5^; *q* = 0.0157) not shown.

**Fig. S8:**
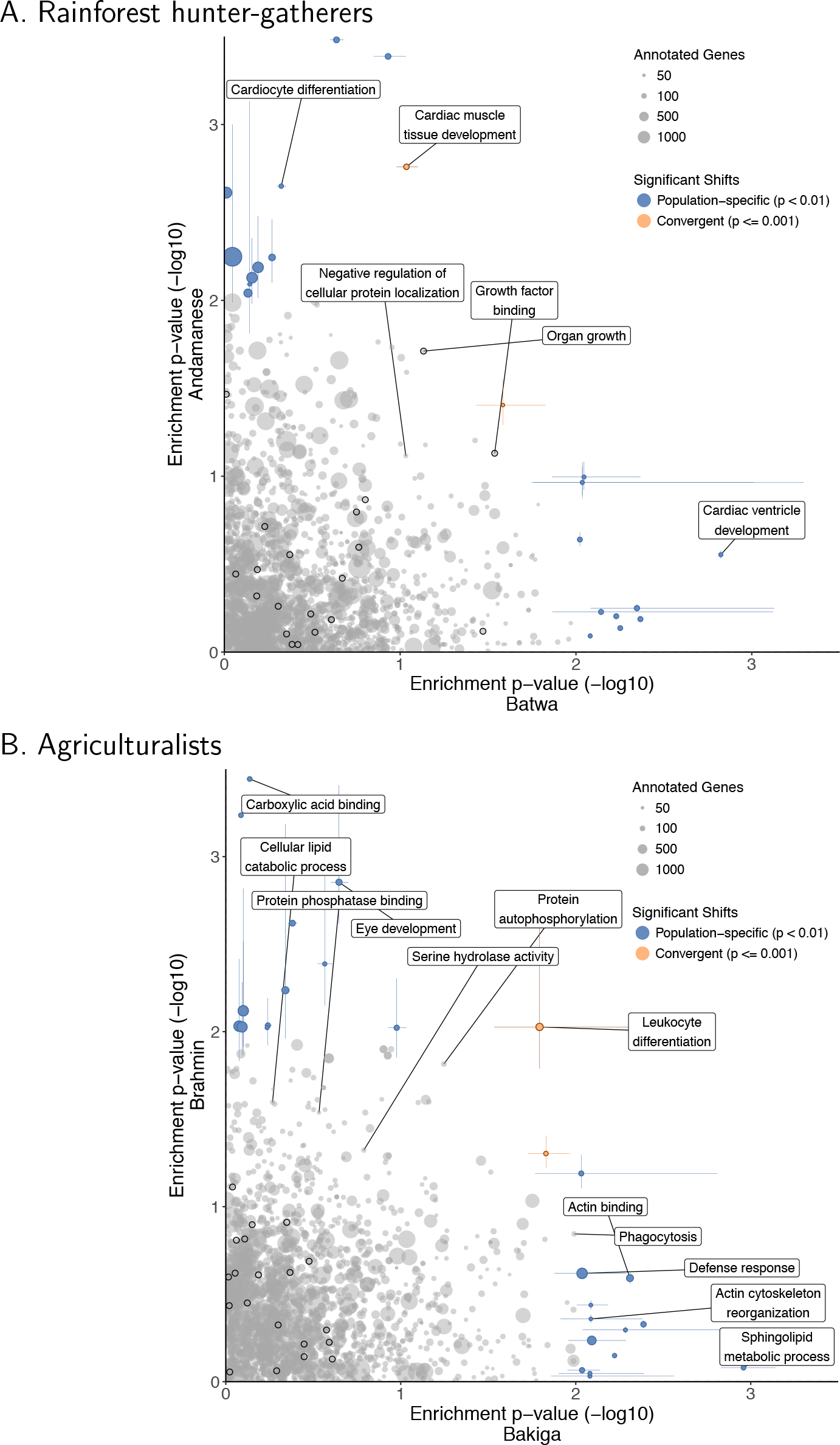
Gene size-corrected polygenic distribution shift test results. After MAF-based correction, Gene Ontology (GO) functional categories’ distribution shift test p-values, indicating a shift in the PBS selection index values for genes, in the Batwa and Andamanese rainforest hunter-gatherers (A) and Bakiga and Brahmin agriculturalist control (B). Results shown for GO biological processes and molecular functions. Point size is scaled to number of annotated genes in category. Terms that are significantly enriched for genes under positive selection (Kolmogorov-Smirnov *p* < 0.01) in either population shown in blue and for both populations convergently (empirical permutation-based *p* < 0.005) shown in orange. Colored lines represent 95% CI for significant categories estimated by bootstrapping genes within pathways. Dark outlines indicate growth-associated terms: the ‘growth’ biological process (GO:0040007) and its descendant terms, or the molecular functions ‘growth factor binding,’ ‘growth factor receptor binding,’ ‘growth hormone receptor activity,’ and ‘growth factor activity’ and their sub-categories.

## 3 Tables

**Table S1:** Gene Ontology (GO) biological processes with evidence of convergent enrichment for strong positive selection in the hunter-gatherer populations, as measured by outlier Population Branch Statistic (PBS) values. No molecular functions were found to be convergently enriched. Joint p-values were computed via a permutation-based method, and those with joint empirical *p* < 0.005 are shown.

| GO Biological Process | Joint $p$ | *Batwa:* | | | | *Andamanese:* | | | |
| --- | --- | --- | --- | --- | --- | --- | --- | --- | --- |
| | | Exp. | Obs. | $p$ | adj. $p$ | Exp. | Obs. | $p$ | adj. $p$ |
| GO:0030326 embryonic limb morphogenesis | 0.000 | 1.47 | 4 | 0.0584 | 1 | 2.01 | 6 | 0.0147 | 0.901 |
| GO:0035107 appendage morphogenesis | 0.000 | 1.69 | 5 | 0.0267 | 1 | 2.27 | 6 | 0.0254 | 0.901 |
| GO:0035108 limb morphogenesis | 0.000 | 1.69 | 5 | 0.0267 | 1 | 2.27 | 6 | 0.0254 | 0.901 |
| GO:0035113 embryonic appendage morphogenesis | 0.000 | 1.47 | 4 | 0.0584 | 1 | 2.01 | 6 | 0.0147 | 0.901 |
| GO:0048736 appendage development | 0.001 | 1.95 | 5 | 0.0448 | 1 | 2.62 | 6 | 0.047 | 1.000 |
| GO:0060173 limb development | 0.001 | 1.95 | 5 | 0.0448 | 1 | 2.62 | 6 | 0.047 | 1.000 |
| GO:0030048 actin filament-based movement | 0.002 | 1.72 | 4 | 0.0927 | 1 | 2.33 | 5 | 0.0834 | 1.000 |
| GO:0048705 skeletal system morphogenesis | 0.002 | 2.20 | 5 | 0.0688 | 1 | 2.95 | 6 | 0.0743 | 1.000 |
| GO:0007034 vacuolar transport | 0.003 | 1.37 | 4 | 0.0470 | 1 | 1.75 | 4 | 0.0968 | 1.000 |

**Table S2:** Gene Ontology (GO) biological processes with evidence of population-specific enrichment for strong positive selection in the hunter-gatherer populations, as measured by outlier Population Branch Statistic (PBS) values. Results with *p* < 0.01 shown.

| GO | | Exp. | Obs. | $p$ | adj. $p$ |
| --- | --- | --- | --- | --- | --- |
| *Batwa RHG - Biological Processes:* |  |  |  |  |  |
| GO:0007517 | muscle organ development | 10 | 4.02 | 0.0069 | 0.708 |
| GO:0045926 | negative regulation of growth | 7 | 2.48 | 0.0118 | 0.708 |
| *Andamanese RHG - Biological Processes:* |  |  |  |  |  |
| GO:0006302 | double-strand break repair | 10 | 3.14 | 0.0011 | 0.171 |
| GO:0070085 | glycosylation | 12 | 4.34 | 0.0013 | 0.171 |
| GO:0000723 | telomere maintenance | 8 | 2.30 | 0.0020 | 0.175 |
| GO:0033365 | protein localization to organelle | 25 | 14.09 | 0.0036 | 0.189 |
| GO:1903827 | regulation of cellular protein localization | 19 | 9.66 | 0.0036 | 0.189 |
| GO:0007569 | cell aging | 6 | 1.62 | 0.0052 | 0.208 |
| GO:0009101 | glycoprotein biosynthetic process | 12 | 5.28 | 0.0065 | 0.208 |
| GO:0034613 | cellular protein localization | 41 | 27.87 | 0.0067 | 0.208 |
| GO:0051179 | localization | 116 | 97.30 | 0.0079 | 0.208 |
| GO:0060249 | anatomical structure homeostasis | 13 | 6.09 | 0.0079 | 0.208 |
| GO:0045596 | negative regulation of cell differentiation | 18 | 9.79 | 0.0090 | 0.215 |
| *Batwa RHG - Molecular Functions:* |  |  |  |  |  |
| GO:0003723 | RNA binding | 26 | 17.66 | 0.028 | 0.732 |
| GO:0043167 | ion binding | 83 | 70.68 | 0.034 | 0.732 |
| *Andamanese RHG - Molecular Functions:* |  |  |  |  |  |
| GO:0043130 | ubiquitin binding | 7 | 1.89 | 0.0026 | 0.177 |
| GO:0008233 | peptidase activity | 17 | 9.58 | 0.0153 | 0.383 |

**Table S3:** Gene Ontology (GO) biological processes (BP) and molecular functions (MF) with evidence of convergent distribution shifts in PBS selection index values in the hunter-gatherer populations. Joint p-values were computed via a permutation-based method, and those with joint empirical *p* < 0.005 are shown.

|  |  |  |  | *Batwa:* |  | *Andamanese:* |  |
| --- | --- | --- | --- | --- | --- | --- | --- |
| GO |  |  | Joint *p* | *p* | adj. *p* | *p* | adj. *p* |
| BP | GO:0035265 | organ growth | 0.001 | 0.0275 | 0.997 | 0.04509 | 1.000 |
|  | GO:0048738 | cardiac muscle tissue development | 0.001 | 0.0461 | 0.997 | 0.00265 | 1.000 |
|  | GO:1903828 | negative regulation of cellular protein localization | 0.001 | 0.0360 | 0.997 | 0.04275 | 1.000 |
|  | GO:0016202 | regulation of striated muscle tissue development | 0.002 | 0.0135 | 0.997 | 0.04406 | 1.000 |
|  | GO:1901861 | regulation of muscle tissue development | 0.002 | 0.0135 | 0.997 | 0.04406 | 1.000 |
|  | GO:0045444 | fat cell differentiation | 0.004 | 0.0573 | 0.997 | 0.04058 | 1.000 |
| MF | GO:0019199 | transmembrane receptor protein kinase activity | 0.000 | 0.027 | 0.817 | 0.0261 | 0.784 |
|  | GO:0019838 | growth factor binding | 0.000 | 0.021 | 0.817 | 0.0269 | 0.784 |
|  | GO:0032559 | adenyl ribonucleotide binding | 0.003 | 0.020 | 0.817 | 0.0579 | 0.877 |
|  | GO:0030554 | adenyl nucleotide binding | 0.004 | 0.017 | 0.817 | 0.0755 | 0.877 |

**Table S4:**
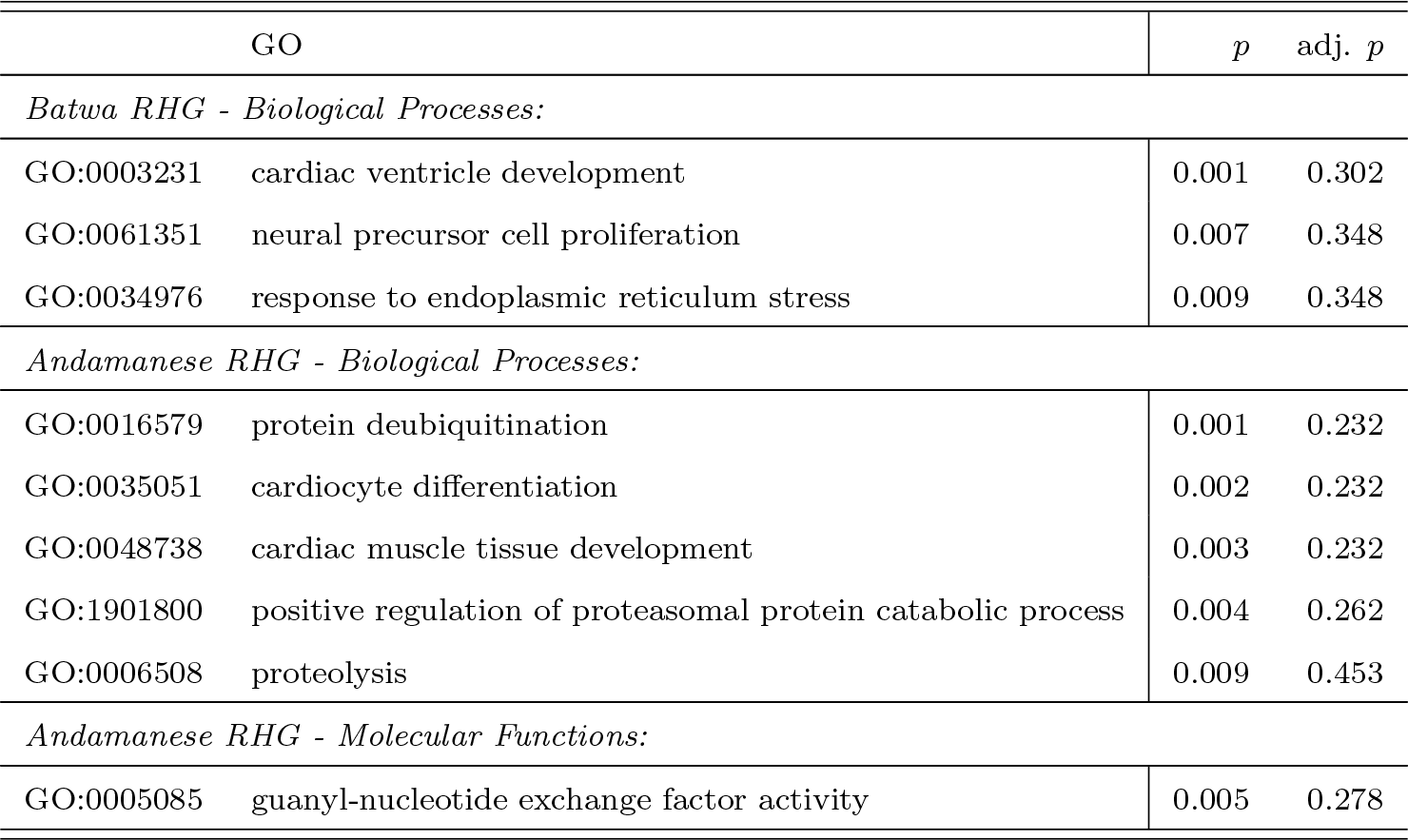
Gene Ontology (GO) biological processes (BP) and molecular functions (MF) with evidence of population-specific distribution shifts in PBS selection index values in the hunter-gatherer populations. No molecular functions were found to be significantly shifted for the Batwa. Results with *p* < 0.01 are shown.

| GO | | $p$ | adj. $p$ |
| --- | --- | --- | --- |
| *Batwa RHG - Biological Processes:* |  |  |  |
| GO:0003231 | cardiac ventricle development | 0.001 | 0.302 |
| GO:0061351 | neural precursor cell proliferation | 0.007 | 0.348 |
| GO:0034976 | response to endoplasmic reticulum stress | 0.009 | 0.348 |
| *Andamanese RHG - Biological Processes:* |  |  |  |
| GO:0016579 | protein deubiquitination | 0.001 | 0.232 |
| GO:0035051 | cardiocyte differentiation | 0.002 | 0.232 |
| GO:0048738 | cardiac muscle tissue development | 0.003 | 0.232 |
| GO:1901800 | positive regulation of proteasomal protein catabolic process | 0.004 | 0.262 |
| GO:0006508 | proteolysis | 0.009 | 0.453 |
| *Andamanese RHG - Molecular Functions:* |  |  |  |
| GO:0005085 | guanyl-nucleotide exchange factor activity | 0.005 | 0.278 |

**Table S5:** After gene size-based correction, Gene Ontology (GO) biological processes and molecular functions with evidence of convergent enrichment for strong positive selection in the hunter-gatherer populations, as measured by outlier Population Branch Statistic (PBS) values. Joint p-values were computed via a permutation-based method, and those with joint empirical *p* < 0.005 are shown.

| GO Biological Process | Joint $p$ | *Batwa:* | | | | *Andamanese:* | | | |
| --- | --- | --- | --- | --- | --- | --- | --- | --- | --- |
| | | Exp. | Obs. | $p$ | adj. $p$ | Exp. | Obs. | $p$ | adj. $p$ |
| GO:0035107 appendage morphogenesis | 0.000 | 1.77 | 5 | 0.0315 | 1 | 2.28 | 6 | 0.0258 | 0.966 |
| GO:0035108 limb morphogenesis | 0.000 | 1.77 | 5 | 0.0315 | 1 | 2.28 | 6 | 0.0258 | 0.966 |
| GO:0048736 appendage development | 0.000 | 2.04 | 5 | 0.0524 | 1 | 2.64 | 6 | 0.0478 | 1.000 |
| GO:0060173 limb development | 0.000 | 2.04 | 5 | 0.0524 | 1 | 2.64 | 6 | 0.0478 | 1.000 |
| GO:1901617 organic hydroxy compound biosynthetic process | 0.000 | 2.38 | 6 | 0.0314 | 1 | 3.19 | 7 | 0.0401 | 0.980 |
| GO:0030326 embryonic limb morphogenesis | 0.001 | 1.53 | 4 | 0.0665 | 1 | 2.02 | 6 | 0.0150 | 0.966 |
| GO:0035113 embryonic appendage morphogenesis | 0.001 | 1.53 | 4 | 0.0665 | 1 | 2.02 | 6 | 0.0150 | 0.966 |
| GO:0030048 actin filament-based movement | 0.002 | 1.80 | 6 | 0.0089 | 1 | 2.34 | 5 | 0.0845 | 1.000 |
| GO:0007034 vacuolar transport | 0.004 | 1.43 | 4 | 0.0537 | 1 | 1.76 | 4 | 0.0979 | 1.000 |
| GO Molecular Function | Joint $p$ | *Batwa:* | | | | *Andamanese:* | | | |
| | | Exp. | Obs. | $p$ | adj. $p$ | Exp. | Obs. | $p$ | adj. $p$ |
| GO:0008514 organic anion transmembrane transporter activity | 0.000 | 2.78 | 7 | 0.0212 | 0.6853 | 3.37 | 7 | 0.0515 | 0.902 |
| GO:0015081 sodium ion transmembrane transporter activity | 0.000 | 2.31 | 7 | 0.0081 | 0.6853 | 2.93 | 6 | 0.0726 | 0.902 |

**Table S6:** After MAF-based correction, Gene Ontology (GO) biological processes and molecular functions with evidence of convergent enrichment for strong positive selection in the hunter-gatherer populations, as measured by outlier Population Branch Statistic (PBS) values. Joint p-values were computed via a permutation-based method, and those with joint empirical *p* < 0.005 are shown.

| GO Biological Process | Joint $p$ | *Batwa:* | | | | *Andamanese:* | | | |
| --- | --- | --- | --- | --- | --- | --- | --- | --- | --- |
| | | Exp. | Obs. | $p$ | adj. $p$ | Exp. | Obs. | $p$ | adj. $p$ |
| GO:0048522 positive regulation of cellular process | 0.000 | 55.57 | 66 | 0.0534 | 1 | 87.93 | 106 | 0.01153 | 0.946 |
| GO:1901617 organic hydroxy compound biosynthetic process | 0.000 | 2.29 | 6 | 0.0267 | 1 | 3.66 | 9 | 0.01069 | 0.946 |
| GO:0006302 double-strand break repair | 0.001 | 2.26 | 5 | 0.0757 | 1 | 3.62 | 8 | 0.02808 | 0.946 |
| GO:0006897 endocytosis | 0.001 | 8.49 | 14 | 0.0447 | 1 | 13.76 | 22 | 0.01948 | 0.946 |
| GO:0048738 cardiac muscle tissue development | 0.001 | 2.52 | 6 | 0.0399 | 1 | 4.01 | 8 | 0.04694 | 0.946 |
| GO:0080135 regulation of cellular response to stress | 0.001 | 7.63 | 14 | 0.0203 | 1 | 11.95 | 20 | 0.01624 | 0.946 |
| GO:0014706 striated muscle tissue development | 0.002 | 4.07 | 8 | 0.0509 | 1 | 6.55 | 14 | 0.00585 | 0.946 |
| GO:0070302 regulation of stress-activated protein kinase signaling cascade | 0.002 | 2.52 | 6 | 0.0399 | 1 | 3.93 | 7 | 0.09815 | 0.946 |
| GO:1901615 organic hydroxy compound metabolic process | 0.002 | 5.70 | 9 | 0.1168 | 1 | 8.87 | 14 | 0.06046 | 0.946 |
| GO:2001020 regulation of response to DNA damage stimulus | 0.002 | 2.09 | 5 | 0.0572 | 1 | 3.35 | 7 | 0.04997 | 0.946 |
| GO:0051592 response to calcium ion | 0.003 | 1.75 | 6 | 0.0079 | 1 | 2.74 | 5 | 0.13789 | 0.946 |
| GO:0015718 monocarboxylic acid transport | 0.004 | 1.63 | 4 | 0.0793 | 1 | 2.39 | 5 | 0.08980 | 0.946 |
| GO:0030048 actin filament-based movement | 0.004 | 1.73 | 4 | 0.0942 | 1 | 2.78 | 7 | 0.02039 | 0.946 |
| GO:0030326 embryonic limb morphogenesis | 0.004 | 1.48 | 4 | 0.0594 | 1 | 2.31 | 5 | 0.08052 | 0.946 |
| GO:0031098 stress-activated protein kinase signaling | 0.004 | 3.15 | 7 | 0.0383 | 1 | 4.97 | 8 | 0.12454 | 0.946 |
| GO:0035113 embryonic appendage morphogenesis | 0.004 | 1.48 | 4 | 0.0594 | 1 | 2.31 | 5 | 0.08052 | 0.946 |
| GO:0060537 muscle tissue development | 0.004 | 4.30 | 8 | 0.0660 | 1 | 6.90 | 14 | 0.00910 | 0.946 |
| GO Molecular Function | Joint $p$ | *Batwa:* | | | | *Andamanese:* | | | |
| | | Exp. | Obs. | $p$ | adj. $p$ | Exp. | Obs. | $p$ | adj. $p$ |
| GO:0008514 organic anion transmembrane transporter activity | 0.002 | 2.68 | 6 | 0.051 | 0.798 | 4.03 | 10 | 0.0068 | 0.959 |
| GO:0005342 organic acid transmembrane transporter activity | 0.003 | 1.97 | 5 | 0.046 | 0.798 | 2.96 | 7 | 0.0282 | 1.000 |
| GO:0015081 sodium ion transmembrane transporter activity | 0.003 | 2.22 | 5 | 0.071 | 0.821 | 3.50 | 7 | 0.0600 | 1.000 |
| GO:0046943 carboxylic acid transmembrane transporter activity | 0.004 | 1.79 | 4 | 0.103 | 0.964 | 2.70 | 7 | 0.0177 | 1.000 |

**Table S7:** After gene size-based correction, Gene Ontology (GO) biological processes with evidence of population-specific enrichment for strong positive selection in the hunter-gatherer populations, as measured by outlier Population Branch Statistic (PBS) values. Results with *p* < 0.01 shown.

| GO | Exp. | Obs. | $p$ | adj. $p$ |
| --- | --- | --- | --- | --- |
| *Batwa RHG - Biological Processes:* |  |  |  |  |
| GO:0007517 muscle organ development | 11 | 4.20 | 0.0032 | 0.5650 |
| GO:1903825 organic acid transmembrane transport | 6 | 1.67 | 0.0061 | 0.5652 |
| GO:0030048 actin filament-based movement | 6 | 1.80 | 0.0061 | 0.5652 |
| *Andamanese RHG - Biological Processes:* |  |  |  |  |
| GO:0006302 double-strand break repair | 10 | 3.16 | 0.0012 | 0.231 |
| GO:0000723 telomere maintenance | 8 | 2.31 | 0.0020 | 0.231 |
| GO:1903827 regulation of cellular protein localization | 19 | 9.70 | 0.0038 | 0.231 |
| GO:0070085 glycosylation | 11 | 4.36 | 0.0042 | 0.231 |
| GO:1900180 regulation of protein localization to nucleus | 10 | 3.77 | 0.0044 | 0.231 |
| GO:0007569 cell aging | 6 | 1.63 | 0.0053 | 0.232 |
| GO:0060249 anatomical structure homeostasis | 13 | 6.12 | 0.0082 | 0.299 |
| GO:0051234 establishment of localization | 99 | 81.24 | 0.0091 | 0.299 |
| *Batwa RHG - Molecular Functions:* |  |  |  |  |
| GO:0015081 sodium ion transmembrane transporter activity | 7 | 2.31 | 0.0081 | 0.33245 |
| *Andamanese RHG - Molecular Functions:* |  |  |  |  |
| GO:0043130 ubiquitin binding | 7 | 1.89 | 0.0026 | 0.177 |

**Table S8:** After MAF-based correction, Gene Ontology (GO) biological processes with evidence of population-specific enrichment for strong positive selection in the hunter-gatherer populations, as measured by outlier Population Branch Statistic (PBS) values. No molecular functions were found to be significantly shifted for the Batwa. Results with *p* < 0.01 shown.

| GO | | Exp. | Obs. | $p$ | adj. $p$ |
| --- | --- | --- | --- | --- | --- |
| *Batwa RHG - Biological Processes:* |  |  |  |  |  |
| GO:0007517 | muscle organ development | 11 | 4.04 | 0.0023 | 0.492 |
| GO:0051592 | response to calcium ion | 6 | 1.75 | 0.0079 | 0.492 |
| *Andamanese RHG - Biological Processes:* |  |  |  |  |  |
| GO:0051179 | localization | 142 | 114.34 | 0.00044 | 0.115 |
| GO:0045596 | negative regulation of cell differentiation | 22 | 11.53 | 0.0026 | 0.266 |
| GO:0071229 | cellular response to acid chemical | 9 | 3.24 | 0.0048 | 0.266 |
| GO:0002460 | adaptive immune response based on somatic recombination... | 11 | 4.47 | 0.0050 | 0.266 |
| GO:0014706 | striated muscle tissue development | 14 | 6.55 | 0.0059 | 0.266 |
| GO:0048584 | positive regulation of response to stimulus | 53 | 38.05 | 0.0067 | 0.266 |
| GO:0016337 | single organismal cell-cell adhesion | 22 | 12.61 | 0.0076 | 0.266 |
| *Andamanese RHG - Molecular Functions:* |  |  |  |  |  |
| GO:0043130 | ubiquitin binding | 7 | 2.24 | 0.0067 | 0.221 |
| GO:0008514 | organic anion transmembrane transporter activity | 10 | 4.03 | 0.0068 | 0.221 |

**Table S9:** After gene size-based correction, Gene Ontology (GO) biological processes (BP) and molecular functions (MF) with evidence of convergent distribution shifts in PBS selection index values in the hunter-gatherer populations. Joint p-values were computed via a permutation-based method, and those with joint empirical *p* < 0.005 are shown.

| GO | | | Joint $p$ | *Batwa:* | | *Andamanese:* | |
| --- | --- | --- | --- | --- | --- | --- | --- |
| | | | | $p$ | adj. $p$ | $p$ | adj. $p$ |
| BP | GO:0016202 | regulation of striated muscle tissue development | 0.000 | 0.0100 | 0.994 | 0.0570 | 1.000 |
|  | GO:1901861 | regulation of muscle tissue development | 0.000 | 0.0100 | 0.994 | 0.0570 | 1.000 |
|  | GO:0045444 | fat cell differentiation | 0.001 | 0.0539 | 0.994 | 0.0517 | 1.000 |
|  | GO:0048634 | regulation of muscle organ development | 0.002 | 0.0101 | 0.994 | 0.0924 | 1.000 |
|  | GO:0035265 | organ growth | 0.003 | 0.0260 | 0.994 | 0.0613 | 1.000 |
|  | GO:0048738 | cardiac muscle tissue development | 0.003 | 0.0646 | 0.994 | 0.0031 | 1.000 |
|  | GO:0051147 | regulation of muscle cell differentiation | 0.003 | 0.0242 | 0.994 | 0.1026 | 1.000 |
|  | GO:1903828 | negative regulation of cellular protein localization | 0.003 | 0.0475 | 0.994 | 0.0405 | 1.000 |
|  | GO:0046434 | organophosphate catabolic process | 0.004 | 0.0154 | 0.994 | 0.0780 | 1.000 |
| MF | GO:0019199 | transmembrane receptor protein kinase activity | 0.000 | 0.021 | 0.698 | 0.0132 | 0.736 |
|  | GO:0019838 | growth factor binding | 0.002 | 0.029 | 0.750 | 0.0285 | 0.736 |
|  | GO:0030554 | adenyl nucleotide binding | 0.002 | 0.014 | 0.698 | 0.0769 | 0.877 |
|  | GO:0032559 | adenyl ribonucleotide binding | 0.002 | 0.017 | 0.698 | 0.0599 | 0.877 |

**Table S10:**
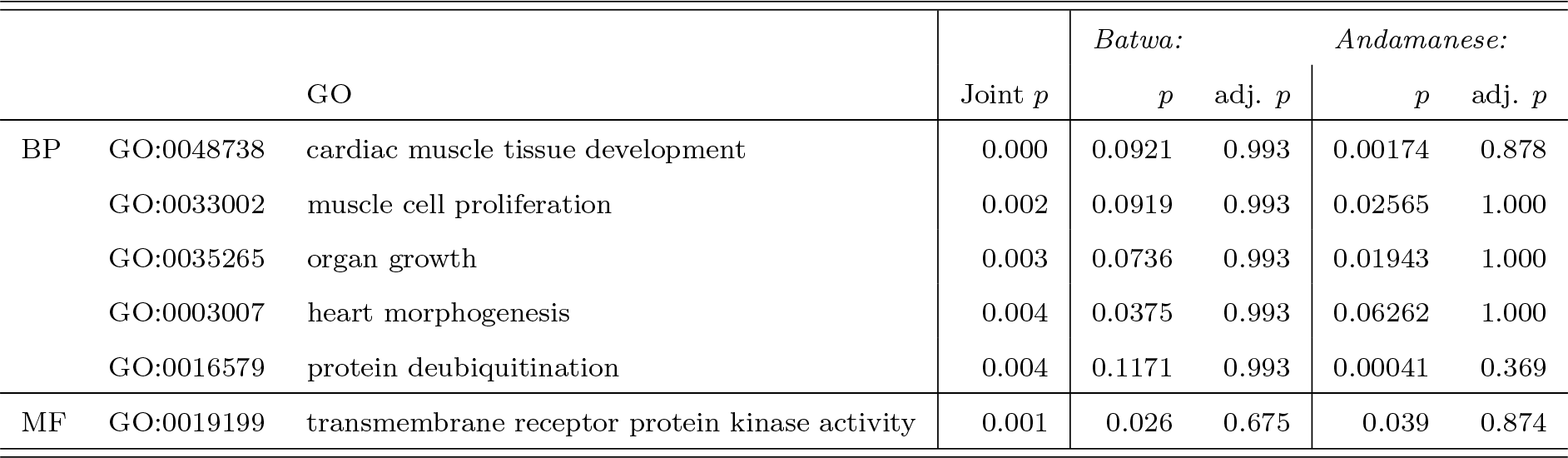
After MAF-based correction, Gene Ontology (GO) biological processes (BP) and molecular functions (MF) with evidence of convergent distribution shifts in PBS selection index values in the hunter-gatherer populations. Joint p-values were computed via a permutation-based method, and those with joint empirical *p* < 0.005 are shown.

**Table S11:** After gene size-based correction, Gene Ontology (GO) biological processes (BP) and molecular functions (MF) with evidence of population-specific distribution shifts in PBS selection index values in the hunter-gatherer populations. No molecular functions were found to be significantly shifted for the Batwa. Results with *p* < 0.01 are shown.

| GO | | $p$ | adj. $p$ |
| --- | --- | --- | --- |
| *Batwa RHG - Biological Processes:* |  |  |  |
| GO:0003231 | cardiac ventricle development | 0.0025 | 0.371 |
| GO:0061351 | neural precursor cell proliferation | 0.0080 | 0.371 |
| *Andamanese RHG - Biological Processes:* |  |  |  |
| GO:0016579 | protein deubiquitination | 0.001 | 0.273 |
| GO:0035051 | cardiocyte differentiation | 0.003 | 0.273 |
| GO:0048738 | cardiac muscle tissue development | 0.003 | 0.273 |
| GO:1901800 | positive regulation of proteasomal protein catabolic process | 0.006 | 0.322 |
| GO:0001936 | regulation of endothelial cell proliferation | 0.006 | 0.322 |
| GO:0006508 | proteolysis | 0.009 | 0.396 |
| *Andamanese RHG - Molecular Functions:* |  |  |  |
| GO:0005085 | guanyl-nucleotide exchange factor activity | 0.0034 | 0.224 |
| GO:0008134 | transcription factor binding | 0.0096 | 0.224 |

**Table S12:** After MAF-based correction, Gene Ontology (GO) biological processes (BP) and molecular functions (MF) with evidence of population-specific distribution shifts in PBS selection index values in the hunter-gatherer populations. No molecular functions were found to be significantly shifted for the Batwa. Results with *p* < 0.01 are shown.

| GO | | $p$ | adj. $p$ |
| --- | --- | --- | --- |
| *Batwa RHG - Biological Processes:* |  |  |  |
| GO:0003231 | cardiac ventricle development | 0.0015 | 0.346 |
| GO:0034284 | response to monosaccharide | 0.0043 | 0.346 |
| GO:0008217 | regulation of blood pressure | 0.0056 | 0.346 |
| GO:0050864 | regulation of B cell activation | 0.0083 | 0.346 |
| GO:0048634 | regulation of muscle organ development | 0.0090 | 0.346 |
| GO:1901861 | regulation of muscle tissue development | 0.0092 | 0.346 |
| *Andamanese RHG - Biological Processes:* |  |  |  |
| GO:0070646 | protein modification by small protein removal | 0.0003 | 0.087 |
| GO:0048738 | cardiac muscle tissue development | 0.0017 | 0.160 |
| GO:0035051 | cardiocyte differentiation | 0.0022 | 0.160 |
| GO:0006508 | proteolysis | 0.0024 | 0.156 |
| GO:0071840 | cellular component organization or biogenesis | 0.0057 | 0.283 |
| GO:0007155 | cell adhesion | 0.0065 | 0.283 |
| GO:0007169 | transmembrane receptor protein tyrosine kinase signaling pathway | 0.0091 | 0.298 |
| *Andamanese RHG - Molecular Functions:* |  |  |  |
| GO:0005085 | guanyl-nucleotide exchange factor activity | 0.0057 | 0.198 |
| GO:0019783 | ubiquitin-like protein-specific protease activity | 0.0081 | 0.198 |

**Table S13:** Comparison of results of two methods for computing empirical test for convergence in strong outlier selection in both the Batwa and Andamanese RHGs. In the original method, genes and PBS selection index values are permuted to create an empirical null distribution. In the modified case, genes and their Gene Ontology (GO) annotations are instead permuted to create the null distribution. Biological processes (BP) with empirical test for convergence *p* < 0.005 in either method shown. No molecular functions were found to be significantly convergently enriched in both RHG populations.

| GO Biological Process | | Original convergence $p$ | Modified convergence $p$ |
| --- | --- | --- | --- |
| GO:0035107 | appendage morphogenesis | 0.000 | 0.000 |
| GO:0035108 | limb morphogenesis | 0.000 | 0.000 |
| GO:0030326 | embryonic limb morphogenesis | 0.000 | 0.002 |
| GO:0035113 | embryonic appendage morphogenesis | 0.000 | 0.002 |
| GO:0048736 | appendage development | 0.001 | 0.003 |
| GO:0060173 | limb development | 0.001 | 0.003 |
| GO:0048705 | skeletal system morphogenesis | 0.002 | 0.003 |
| GO:0048522 | positive regulation of cellular process | 0.006 | 0.003 |
| GO:0080135 | regulation of cellular response to stress | 0.018 | 0.004 |
| GO:0030048 | actin filament-based movement | 0.002 | 0.006 |
| GO:0007034 | vacuolar transport | 0.003 | - |

**Table S14:**
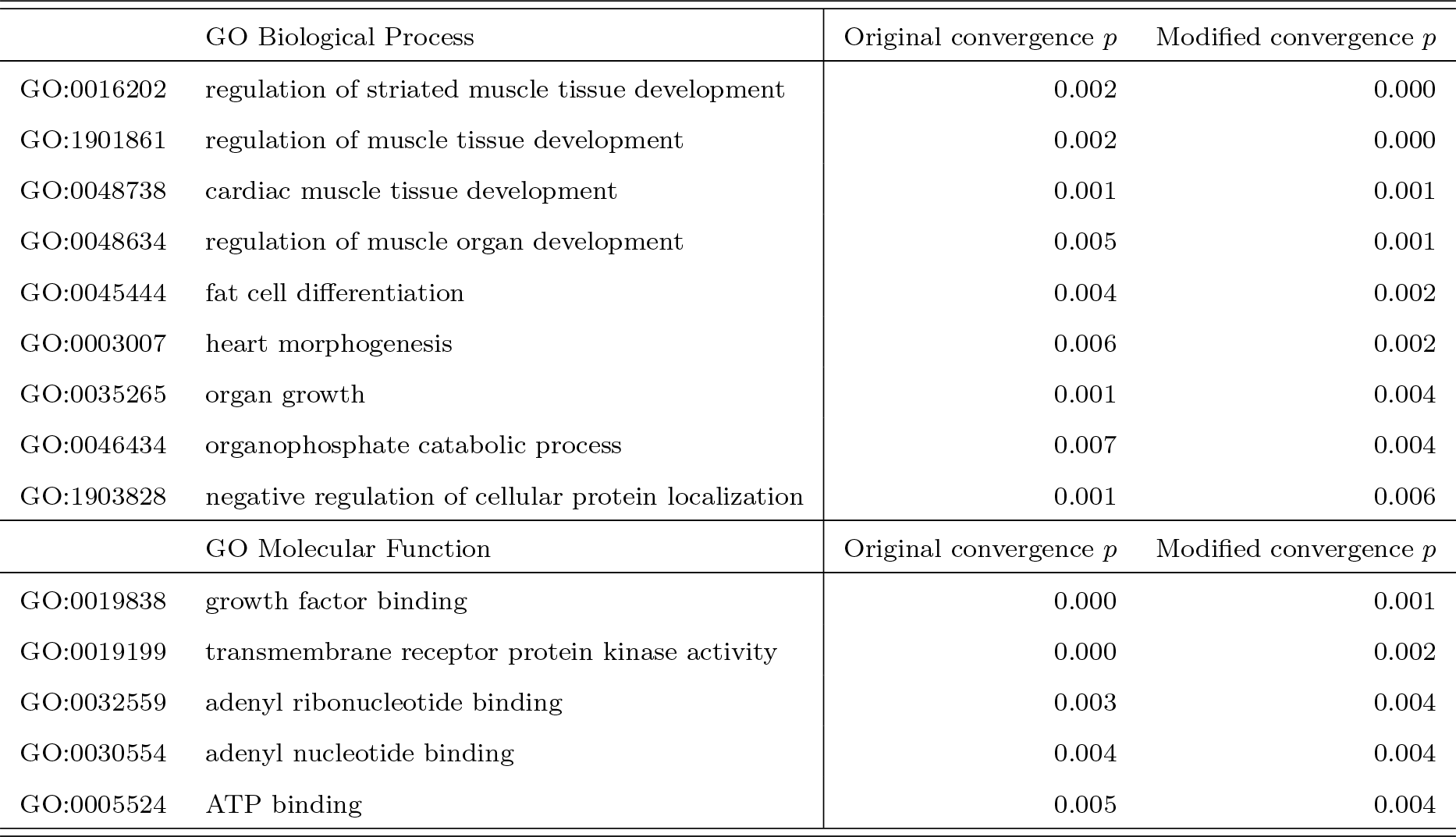
Comparison of results of two methods for computing empirical test for convergence in PBS selection index shift in both the Batwa and Andamanese RHGs. In the original method, genes and PBS selection index values are permuted to create an empirical null distribution. In the modified case, genes and their Gene Ontology (GO) annotations are instead permuted to create the null distribution. Biological processes (BP) and molecular functions (MF) with empirical test for convergence *p* < 0.005 in either method shown.

